# Interaction of HelQ helicase with RPA modulates RPA-DNA binding and stimulates HelQ to unwind DNA through a protein roadblock

**DOI:** 10.1101/511758

**Authors:** Sarah J. Northall, Tabitha Jenkins, Denis Ptchelkine, Vincenzo Taresco, Christopher D. O. Cooper, Panos Soultanas, Edward L. Bolt

## Abstract

Cells reactivate compromised DNA replication forks using enzymes that include DNA helicases for separating DNA strands and remodelling protein-DNA complexes. HelQ helicase promotes replication-coupled DNA repair in mammals in a network of interactions with other proteins. We report newly identified HelQ helicase activities, when acting alone and when interacting with RPA. HelQ helicase was strongly inhibited by a DNA-protein barrier (BamHI^E111A^), and by an abasic site in the translocating DNA strand. Interaction of HelQ with RPA activated DNA unwinding through the protein barrier, but not through the abasic site. Activation was lost when RPA was replaced with bacterial SSB or DNA binding-defective RPA, RPA^ARO1^. We observed stable HelQ-RPA-DNA ternary complex formation, and present evidence that an intrinsically disordered N-terminal region of HelQ (N-HelQ) interacts with RPA, destabilising RPA-DNA binding. Additionally, SEC-MALS showed that HelQ multimers are converted into catalytically active dimers when ATP-Mg^2+^ is bound. HelQ and RPA are proposed to jointly promote replication fork recovery by helicase-catalysed displacement of DNA-bound proteins, after HelQ gains access to ssDNA through its N-terminal domain interaction with RPA.

## INTRODUCTION

DNA replication is essential for the cell cycle of prokaryotes and eukaryotes, and relies on DNA repair and genome stability enzymes to ensure that genome duplication is completed with high fidelity. Encounters of the replisome, the multi-enzyme machine that synthesises new DNA, with chemical damage to DNA or DNA-bound protein complexes stresses DNA replication causing it to halt or collapse and provoking genome instability (1-3). To overcome these problems cells utilize multiple “skipping” and “restart” mechanisms to resume DNA replication according to the disposition of the problem encountered, recently reviewed (4).

DNA replication thwarted by DNA strand breaks can be restarted by homologous recombination, dependent on a strand invasion reaction catalysed by recombinases and their mediator proteins (5-9). Recombination events subsequent to strand invasion take a variety of guises leading to renewed DNA synthesis and reloading of the replisome (10). Replisome blocking lesions on leading or lagging strand templates of the replication fork in *E. coli* can be overcome by “skipping” the problem and re-priming replication downstream of the lesion leaving, ssDNA gaps for repair (11, 12). In eukaryotes a lagging strand block or lesion is overcome by re-priming of DNA synthesis using Pol α (13) but mechanisms to resume blocked leading strand synthesis are less clear. Leading strand synthesis re-primed beyond a DNA lesion (e.g. a cyclobutane pyrimidine dimer) is inefficient, is inhibited by the single strand DNA (ssDNA) binding protein RPA, and may primarily occur by processes other than those catalysed by Pol α (13).

Accumulation of ssDNA is a prominent physical characteristic of stressed DNA replication triggered by factors that slow or halt DNA polymerase activity, uncoupling it from the replicative helicase activity (3). Binding of RPA to ssDNA triggers ATR and ATRIP proteins to co-ordinate DNA repair responses that remove DNA damage or blockages and reactivate productive DNA replication (7, 14, 15). DNA intra-strand mono-adducts and inter-strand crosslinks (ICLs) block DNA replication, causing polymerase-helicase uncoupling (16). ICLs arise naturally from DNA reacting with endogenous aldehydes and other nucleophiles (17, 18), and from treatment of cells with chemotherapeutics (19, 20). Replication and repair past ICLs in mammals requires proteins of the Fanconi anaemia (FA) complementation group (21, 22) and non-FA enzymes including HelQ helicase (originally identified as *mus301*) (23-26) and its homologous helicase-polymerase PolQ (*mus308*) (27-29). Loss of HelQ in ovarian cells is associated with cancer (25) and expression of HelQ is a marker for resistance to chemotherapy in ovarian cancers (30, 31). HelQ localises to replication forks when cells are stressed by camptothecin (32), and physically interacts with ATR, the FA family proteins FancD2/FancI, RPA70 single strand DNA (ssDNA) binding protein, and with a Rad51 paralogue complex comprising Rad51B, Rad51C, Rad51D and XRCC2 (BCDX2)(23, 25). Interaction between *C. elegans* HelQ and Rad51 recombinase dissociates Rad51 from duplex DNA independently of HelQ ATPase or helicase activity (33), which may promote DNA replication fork repair and reactivation without recourse to later stages of homologous recombination (34, 35).

HelQ is a ssDNA-dependent ATPase that translocates ssDNA with 3’ to 5’ polarity (26) and unwinds DNA duplexes within a variety of forked and other branched DNA substrates (32). Sequence orthologues and homologues of HelQ are present throughout mammals, protozoans and plants. Activities of archaeal homologues (Hel308 or Hjm) have been described in several species (36-41).

Here we investigated ATP-dependent DNA helicase activities of human HelQ, in particular its association with ssDNA binding protein RPA. We report that interaction of HelQ with RPA stimulates DNA unwinding through a protein-DNA barrier, and that interaction of RPA with a non-catalytic, non-DNA binding fragment of HelQ modulates RPA-DNA binding. The data suggests that assembly of an RPA-HelQ complex on ssDNA recruits HelQ to sites of stressed replication, and generates the means to remove other DNA-bound proteins by unwinding DNA as an RPA-HelQ complex.

## RESULTS

### HelQ helicase is blocked by chemical modifications to DNA and by protein-DNA barriers

Purified human HelQ protein (Supplementary Figure S1A) is a single-stranded DNA (ssDNA) – dependent ATPase that unwinds model forked DNA by loading onto 3’ tailed ssDNA and translocating with 3’ to 5’ polarity (26), summarised in Supplementary Figures S1B and S1C using protein purified for this work. We set out to learn more about how ATP-dependent translocation and helicase activities of human HelQ might promote DNA replication coupled repair. For ssDNA translocation we began by incorporating single chemical modifications into the 3’ to 5’ tracking strand of a model DNA fork, as a probe for measuring changes in HelQ helicase activity (Figure 1A). In our experimental conditions HelQ (160nM) unwound maximally 60% of unmodified forked DNA (Fork-A, 25 nM) as a function of time, reduced to 20% unwinding by incorporation of either a methyl-phosphonate or phosphorothioate DNA backbone modifications (Figure 1A, respectively Fork-Me and Fork-S), or to 7% unwinding of DNA fork with a single abasic site (Figure 1A, Fork-AP). Inhibition of HelQ DNA helicase activity by the same chemical modifications was also observed in assays titrating HelQ from 0 – 160 nM (Supplementary Figure S2A). Binding of HelQ to each chemically modified fork in EMSAs was not significantly impaired compared to unmodified fork (Supplementary Figure S2B) indicating that blocked DNA translocation is the most likely cause of reduced DNA unwinding. Therefore inhibition of HelQ by modifications to DNA backbone and bases suggests that efficient translocation by HelQ requires electrostatic interaction with the DNA backbone and additional interactions with individual bases.

**Figure 1.**
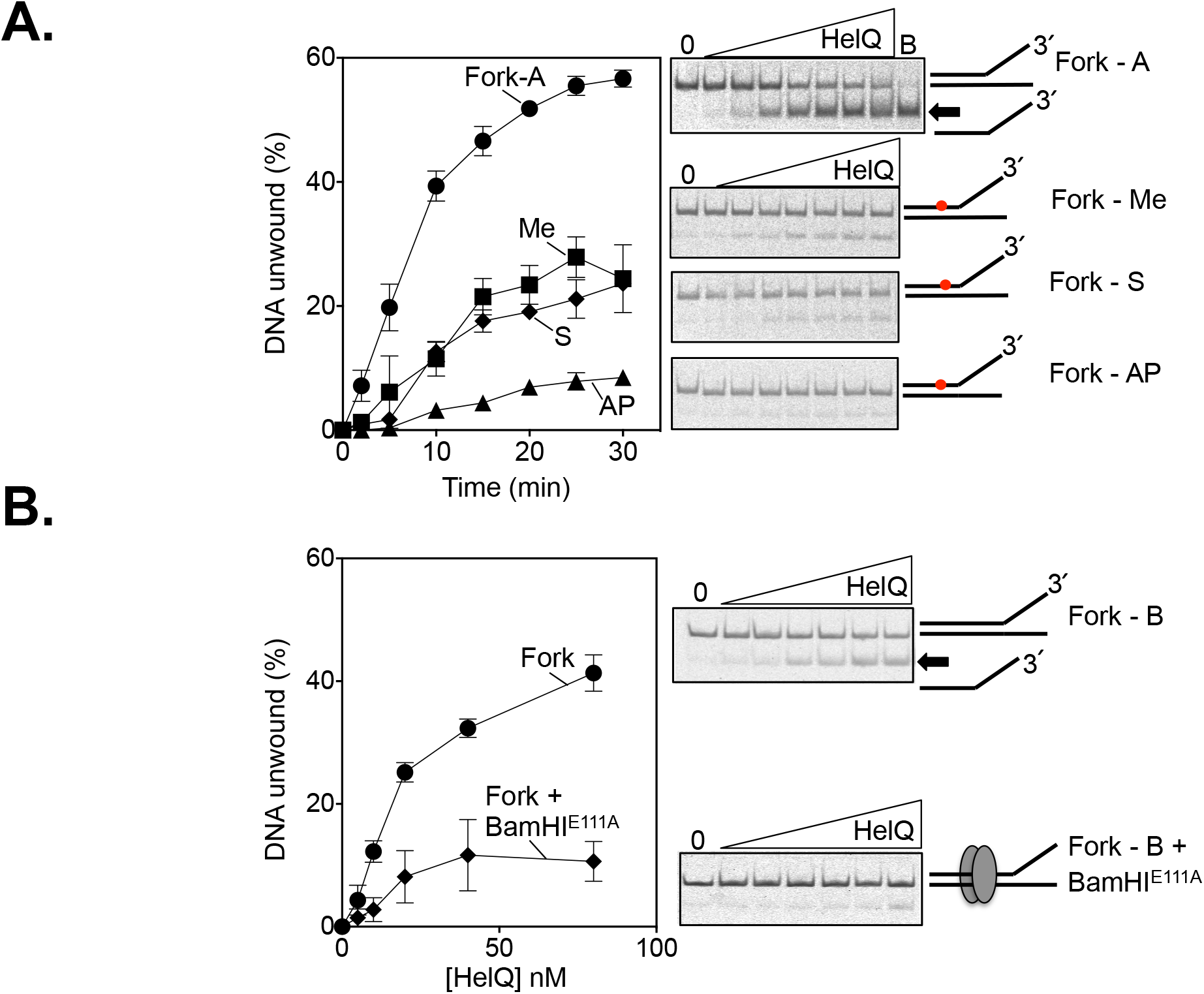
DNA chemical and protein barriers impede translocation by human HelQ. A. DNA helicase activity of human HelQ (150 nM) measured as a function of time when incubated with the DNA forks (25 nM) shown in Supplementary Figure S6. Forks were either unmodified (Fork-A) or contained a chemical modification at a single position on the HelQ tracking DNA strand; methyl-phosphonate (Me), phosphorothioate (S) or an abasic site (AP). The graph shows mean values of three experiments for each fork and error bars of standard deviation from the mean. Representative gel panels corresponding to one assay for each fork are shown to the right of the graph, indicating the chemical modification with a red dot and the unwound product with an arrow, and boiling of the fork into ssDNA as B. B. DNA helicase activity of human HelQ (0, 5, 10, 20 40 and 80 nM) measured on a model DNA fork (25 nM) bound with BamHI^E111A^ (80 nM, diamonds) or unbound (circles), with representative gels shown to the right of the graph illustrated as in (A).

DNA unwinding by HelQ was also strongly inhibited when confronted by a protein-DNA blockage, nucleolytically inactive BamHI (BamHI^E111A^) (Figure 1B). Similar inhibition of DNA unwinding has been reported for human FancJ, RecQ1 and WRN helicases (42). In these assays we used a DNA fork (Fork-B, 25 nM) containing hexa-nucleotide cognate sequence for BamHI DNA binding. Fork-B was 100% bound by BamHI^E111A^ added to 80 or 160 nM in a stable complex seen in EMSAs (Supplementary Figure S2C). Pre-incubation of BamHI^E111A^ with Fork-B reduced HelQ unwinding activity from a mean of 52% to 9 – 20% using BamHI at 80 or 160 nM (Figure 1B), conditions when BamHI^E111A^ bound all fork DNA substrate. HelQ alone is weakly active at translocating through this nucleoprotein complex.

### HelQ and RPA, but not RPA^Aro1^ or *E. coli* SSB, jointly unwind DNA through a protein barrier

HelQ forms a complex with the RPA70 (RPA1), the ssDNA binding subunit of heterotrimeric RPA complex (43, 44), when pulled-down from human cells (23), and co-localises with RPA foci in camptothecin treated human U20S cells (32). Consistent with this, RPA interacted with purified human HelQ immobilised on streptactin beads in the absence of DNA (Figure 2A). We also observed evidence for a stable ternary complex between heterotrimeric RPA, HelQ and DNA in EMSAs (Figure 2B). DNA EMSAs of either HelQ or RPA heterotrimer alone mixed with fork-A gave well-defined protein-DNA complexes, summarised in Figure 2B lanes 2 - 5. Addition of HelQ to fork-A pre-bound with RPA generated a clearly defined ternary “super-shifted” complex (Figure 2B, lanes 6 and 7 marked by an asterisk). Such a super-shifted complex was not observed when the *E. coli* ssDNA binding protein SSB was substituted for RPA (Supplementary Figure S3A), consistent with formation of a specific RPA - HelQ complex rather than generic co-incident ssDNA binding by both proteins.

**Figure 2.**
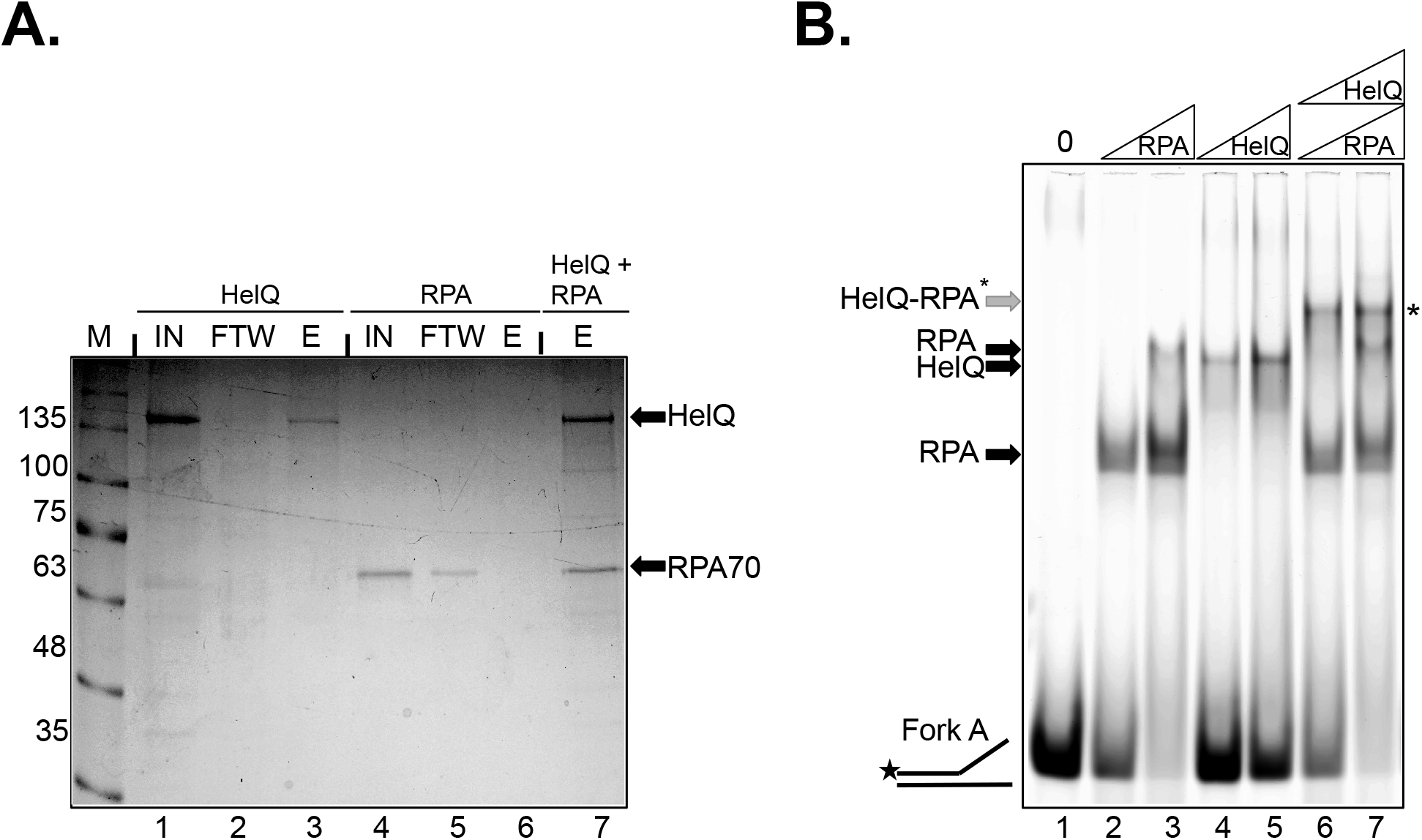
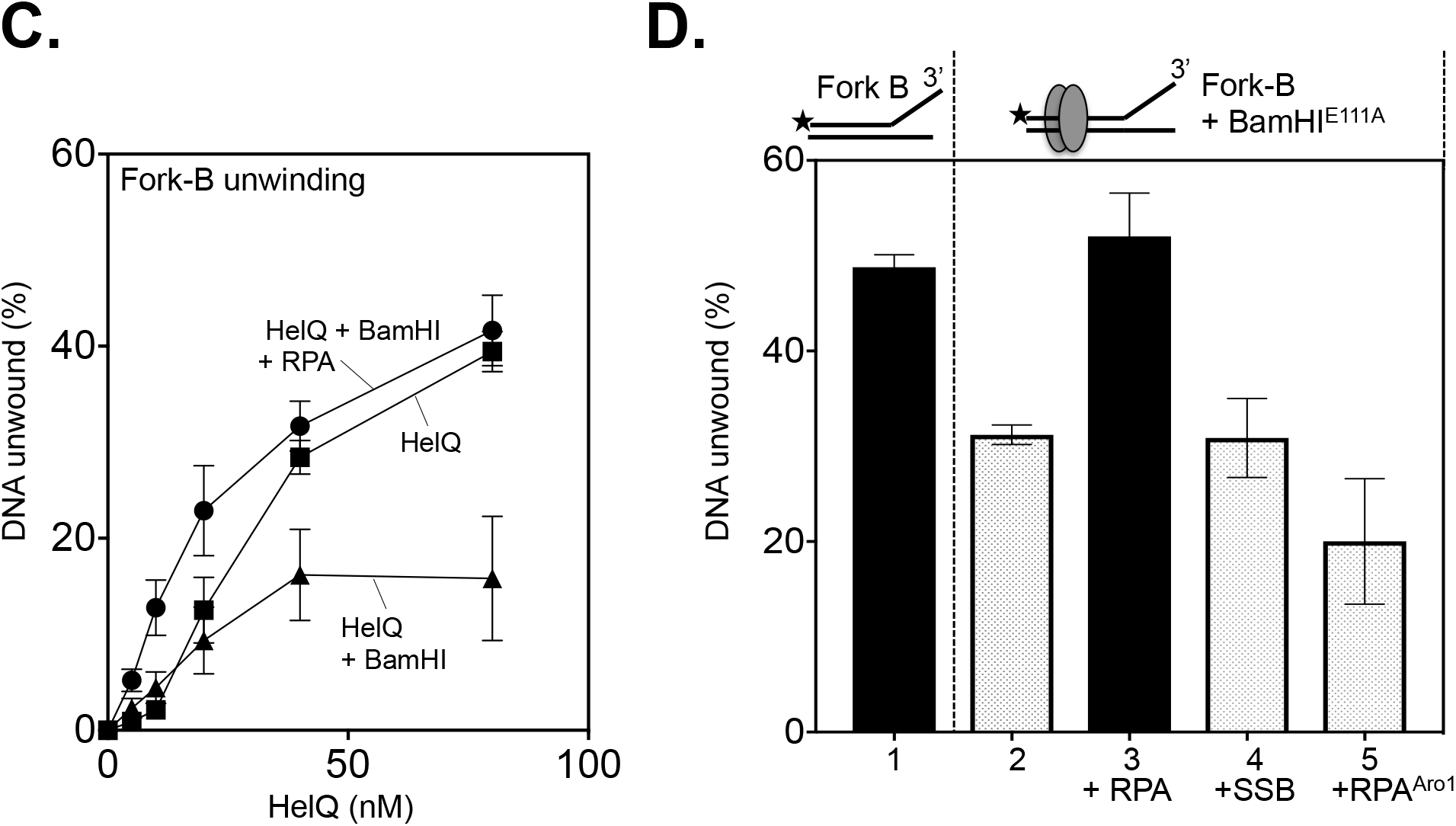
Physical and functional Interaction of HelQ with RPA. A. Coomassie stained 10 % acrylamide SDS-PAGE gel summarising pull-down of RPA70 by streptactin-tagged HelQ. Lanes show protein “input” (IN) mixed with streptactin beads, unbound protein as “flow-through-wash” (FTW), and bound protein eluted from beads (E). Input HelQ alone bound to beads but RPA70 alone did not (lanes 1 - 3 and 4 - 6), both as expected. Pre-mixing RPA70 with HelQ recovered RPA70 into bound fraction E (lane 7). B. EMSA of fork-A DNA (25 nM) bound by RPA (lanes 2 and 3, 5 nM, 20 nM) or HelQ (lanes 4 and 5, 5 or 20 nM), and when RPA was pre-bound to DNA prior to addition of HelQ (lanes 6 and 7). The asterisk (*) indicates ternary HelQ-RPA complex. C. Addition of RPA (5 nM) to HelQ (80 nM) and DNA fork-B bound by BamHI^E111A^ stimulates HelQ to unwind the fork efficiently as when BamHI^E111A^ is not bound, as indicated. The graph panel shows data from three experiments with error bars of deviation from the mean. D. Unwinding of DNA fork-B (25 nM) by HelQ (80 nM) (histogram 1) that is blocked by BamHI^E111A^ (histogram 2) was stimulated by addition of RPA (5 nM, histogram 3) but not by addition of *E. coli* SSB or by human RPA^Aro1^ mutant, respectively 7.5 nM and 5 nM (histograms 4 and 5).

Previous studies showed that human RPA stimulates HelQ to unwind duplex DNA and that this is not due to RPA preventing re-annealing of dissociated DNA strands (26, 32). We observed that RPA stimulated HelQ to overcome the BamHI^E111A^ blockage to helicase activity (Figure 2C and 2D and supplementary data). HelQ (160 nM) combined with RPA (5 nM) restored maximal fork DNA unwinding, a six-fold increase, to levels observed in the absence of BamHI^E111A^ (Figure 2C). Addition to HelQ helicase reactions of *E. coli* SSB or a DNA binding defective RPA mutant protein (RPA^Aro1^) did not stimulate HelQ to unwind DNA through BamHI^E111A^ (Figure 2D and Supplementary Figure S3B). RPA^Aro1^ protein has reduced ssDNA binding caused by mutations of phenylalanine and tryptophan residues in DBD-A and DBD-B domains of RPA70 (45). A “super-shifted” EMSA ternary complex was observed between RPA^Aro1^-HelQ and fork DNA but was significantly altered compared to wild type RPA (Supplementary Figure S3C), further evidence for specific physical interaction between HelQ and RPA. Further control assays confirmed that DNA binding by RPA alone was unable to displace BamHI^E111A^, that stimulation of HelQ helicase activity by RPA was ATP-dependent (Supplementary Figure S3D), and that RPA did not stimulate HelQ ATPase activity (Supplementary Figure S3E). We also assessed if RPA stimulates HelQ helicase to dissociate DNA within a G4 Quadruplex after loading onto a 15-nucleotide polyA 3’ ssDNA tail, a substrate that is unwound weakly by HelQ alone compared to bacterial RecQ (Supplementary Figure S3F). RPA had no stimulatory effect on HelQ unwinding G4 DNA, or on unwinding through the abasic site shown in Figure 1 (Supplementary Figure S3G). These observations indicate that a physical interaction between RPA and HelQ mediates helicase unwinding through a protein-DNA complex.

### A non-catalytic N-HelQ domain interacts with RPA and modulates RPA-DNA binding

We next used purified fragments of HelQ protein to better understand how HelQ and RPA interact. We focussed on using purified HelQ winged helix domain (WHD)(41), based on known interactions of RPA with WHDs in other helicases, and a purified predicted N-terminal domain of HelQ (N-HelQ). N-HelQ comprises 400 - 500 amino acids of unknown function lacking significant primary sequence conservation with other proteins, but we identified predicted structural similarity between N-HelQ and a PWI-like fold that in yeast Brr2 helicase is non-catalytic and is needed for protein interactions (46) (Figure 3A and 3B and Supplementary Figure S4A). HelQ WHD was purified as described previously (41), and a 417-residue (47 kDa) N-HelQ protein was purified as a proteolytic product of full-length HelQ or by recombinant expression in *E. coli* (Figure 3C and Supplementary Figure S4B). WHD or N-HelQ was incubated with fork-A DNA alone or pre-bound with RPA, and protein-DNA complex formation was assessed in EMSAs similar to those used for full-length HelQ. RPA (12.5 nM) bound to 100% of fork DNA (Figure 3D, lane 2) and WHD HelQ (200 nM) bound unstably to DNA as expected (41)(lane 3) but did not generate a convincing ternary complex with RPA (lane 4). N-HelQ (200 nM) did not bind to fork DNA (Figure 3D lane 5, and see Figure 3E) but gave a clearly defined “super-shifted” ternary complex when RPA was present (Figure 3D lane 6, arrowed asterisk). Appearance of the ternary complex corresponded to liberation of free-fork DNA from RPA binding (lane 6, also arrowed asterisk). This was especially apparent as a function of increasing concentration of N-HelQ (0 – 400 nM) when titrated into RPA pre-bound to fork DNA, summarised in Figure 3E and 3F. Initial appearance of a clearly defined new EMSA complex from adding N-HelQ to DNA pre-bound with RPA (asterisked in lanes 3 – 5) subsides, corresponding to liberation of free fork DNA and reduced RPA-DNA complex (asterisked in lanes 5 - 7). Addition of a wide range of N-HelQ concentrations to RPA-DNA reactions was observed to reduce RPA-DNA binding from 100% to 25% (Figure 3F). N-HelQ did not give any observable DNA binding at any concentration used (Figure 3E lanes 8 – 13 and Supplementary Figure S4B). Apparent destabilisation of RPA-DNA binding by N-HelQ was also observed when RPA was bound to 50-nucleotide ssDNA instead of fork, and when magnesium was included in buffers and EMSA gels instead of EDTA (Supplementary Figure S4C). We propose that interaction of the N-terminal region of HelQ with RPA has evolved to modulate RPA-DNA binding in a way that promotes HelQ helicase activity.

**Figure 3.**
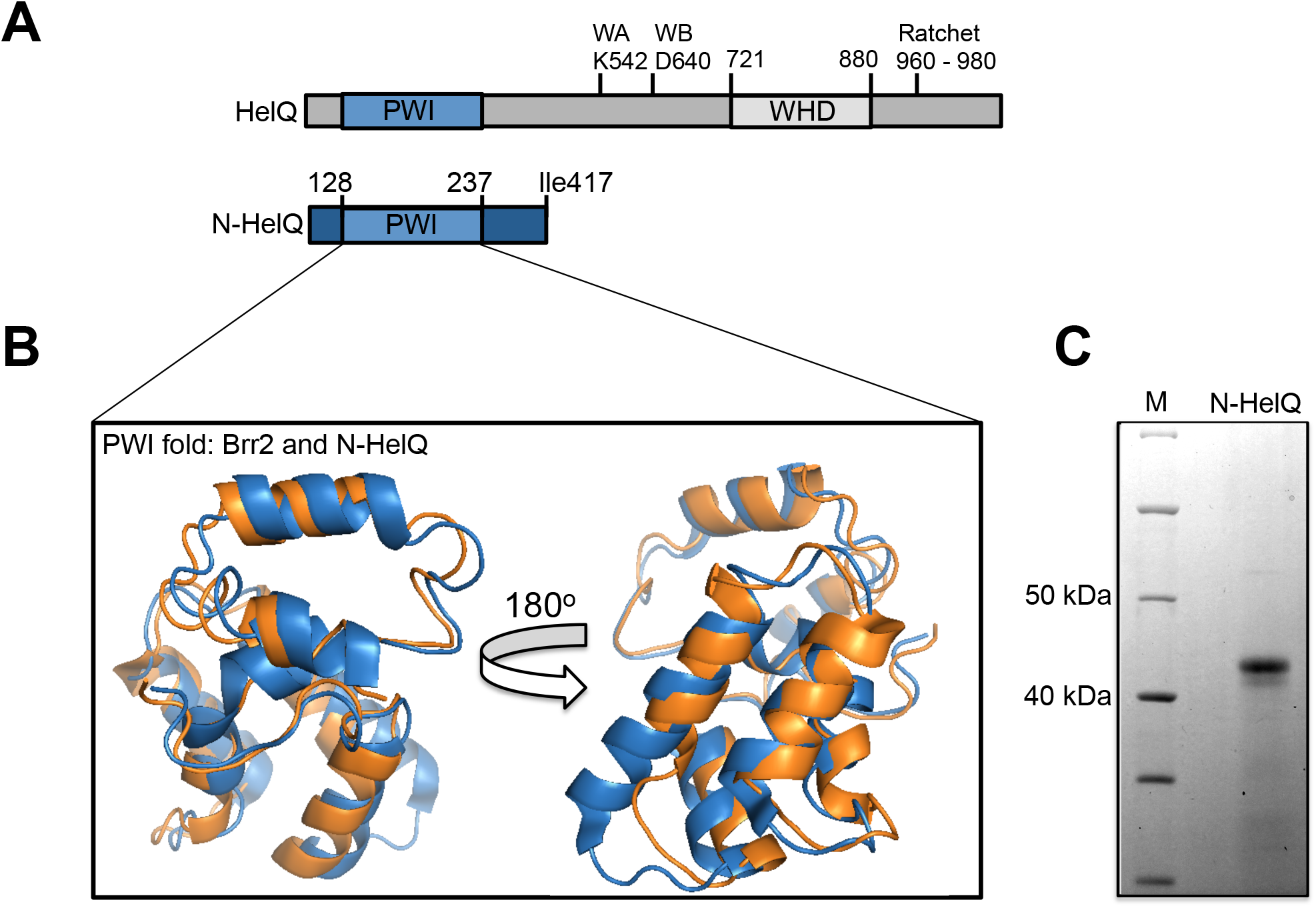
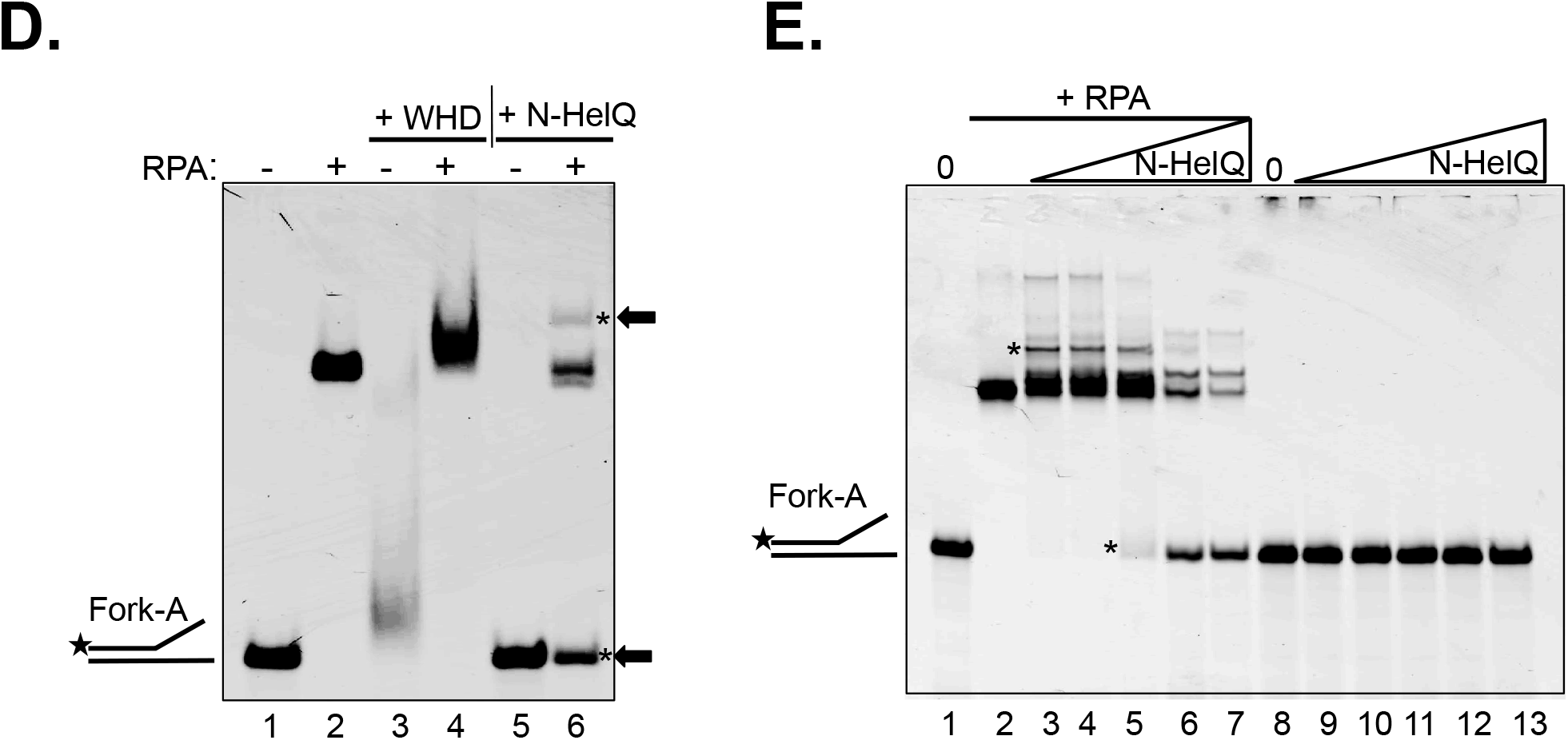
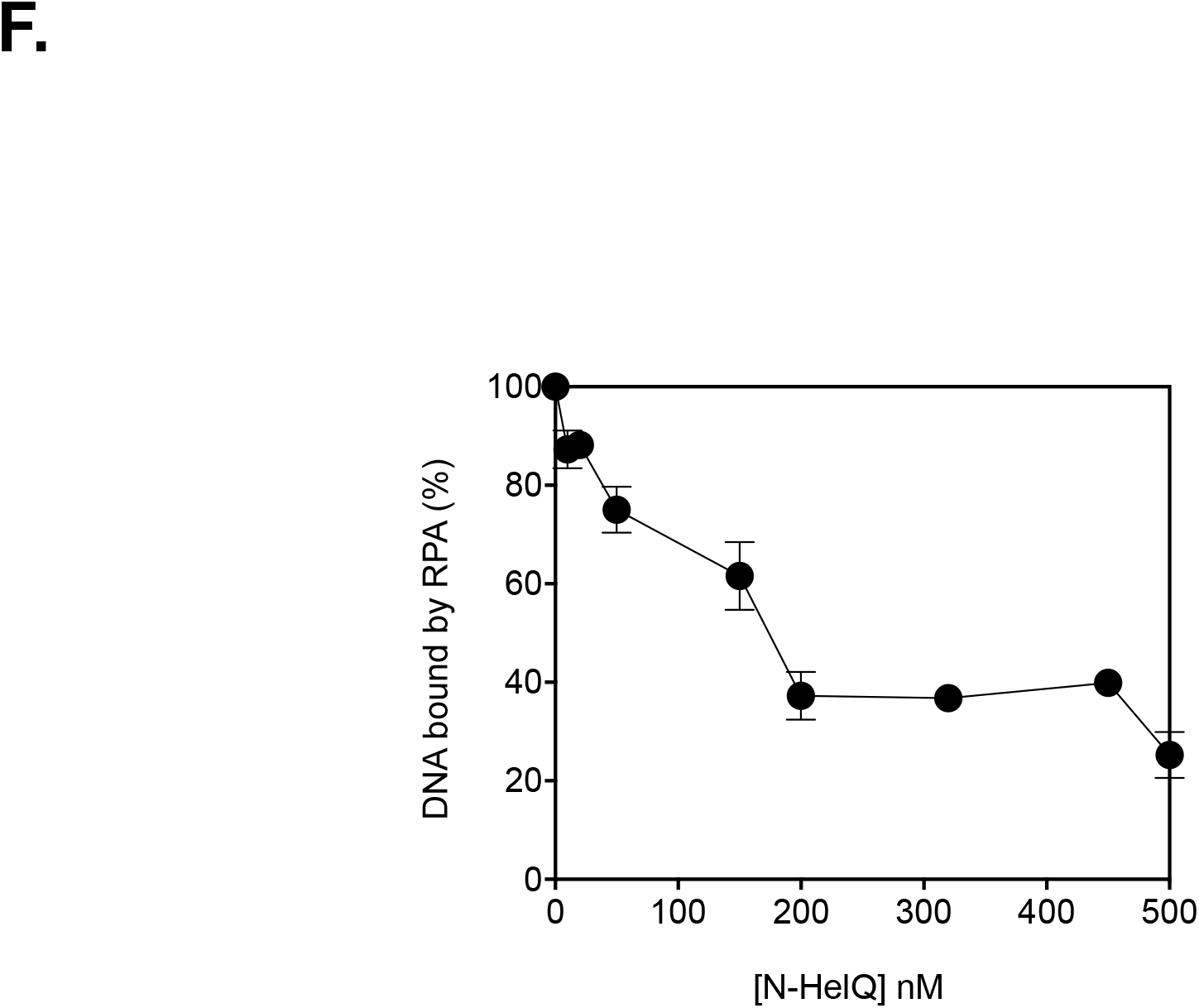
Interaction of N-HelQ and RPA proteins. A. Scheme showing full-length HelQ protein with the non-catalytic PWI-fold located within a 417 amino acid fragment of HelQ that was purified in this work. Helicase motifs are located outside of N-HelQ, including as indicated the Walker A and B sequences (WA, WB) and the helicase “ratchet” helix predicted to be preserved from atomic structures of homologous archaeal Hel308. The position of the winged helix domain (WHD) is also highlighted in full-length HelQ. B. The S. *cerevisiae* Brr2 PWI fold taken from PDB accession 5DCA (46) is shown in blue superimposed with the PHYRSE and DALI modelled human HelQ PWI-like region within N-HelQ (orange). C. Coomassie stained SDS-PAGE gel of purified human N-HelQ protein fragment. See also Supplementary Figure S4B. D. EMSA summarising DNA binding of fork-A (25 nM) by RPA heterotrimer (12.5 nM, lane 2) in the presence and absence of purified HelQ winged helix domain protein fragment (WHD, 200 nM, lanes 3 – 4) or purified N-HelQ fragment (200 nM, lanes 5 – 6). Asterisks and arrows are added to highlight new protein DNA complex formation by mixing RPA with N-HelQ, corresponding to liberation of free fork DNA, despite N-HelQ alone not binding to DNA. E. EMSA showing binding of RPA heterotrimer (12.5 nM) to fork-A DNA (25nM, lane 2) and RPA binding when N-HelQ protein was added to 25, 50, 100, 200 and 300 nM (lanes 3 – 7). Lanes 9 – 13 show that N-HelQ protein added alone to fork-A at the same concentrations show no DNA binding complex. F. Modulation of RPA – fork-A DNA binding (12.5 nM RPA heterotrimer; 25 nM DNA) by addition of N-HelQ, quantified using titration of N-HelQ from 0 – 500 nM into RPA - DNA EMSA reactions. The graph shows reactions in duplicate with error bars for standard error from the mean.

### HelQ dimerises on ATP binding

Protein expression constructs used in this work encoded HelQ monomers of 141 KDa, which includes tandem N-terminal streptactin and (His-)_6_ affinity tags (Figure S1A). Analytical gel filtration (AGF) of purified HelQ protein gave a major HelQ peak corresponding to high molecular mass species (>670 kDa, peak 1 in Figure 4A). This agrees with the study first characterising purified human HelQ helicase, which identified a 600-kDa HelQ protein peak by gel filtration in buffers in the absence of ATP-Mg^2+^ (26). This was confirmed using SEC-MALS of HelQ in the same buffer, which gave a protein mass of 598 kDa (Figure 4B part i). However, addition of ATP-Mg^2+^ to buffers resulted in a pronounced shift in HelQ retention during AGF (peak 2 in Figure 4A) that in SEC-MALS gave a mass of 265 kDa (Figure 4B part ii). A similar mass (240 kDa) was observed when SEC-MALS was repeated with addition of ssDNA to HelQ-ATP-Mg^2+^ (supplementary Figure S5). HelQ in fractions spanning the 265kDa protein peak (peak 2) in ATP-Mg^2+^ bound to fork-A DNA and unwound it (Figure 4C), consistent with HelQ dimers being an active form of the helicase.

**Figure 4.**
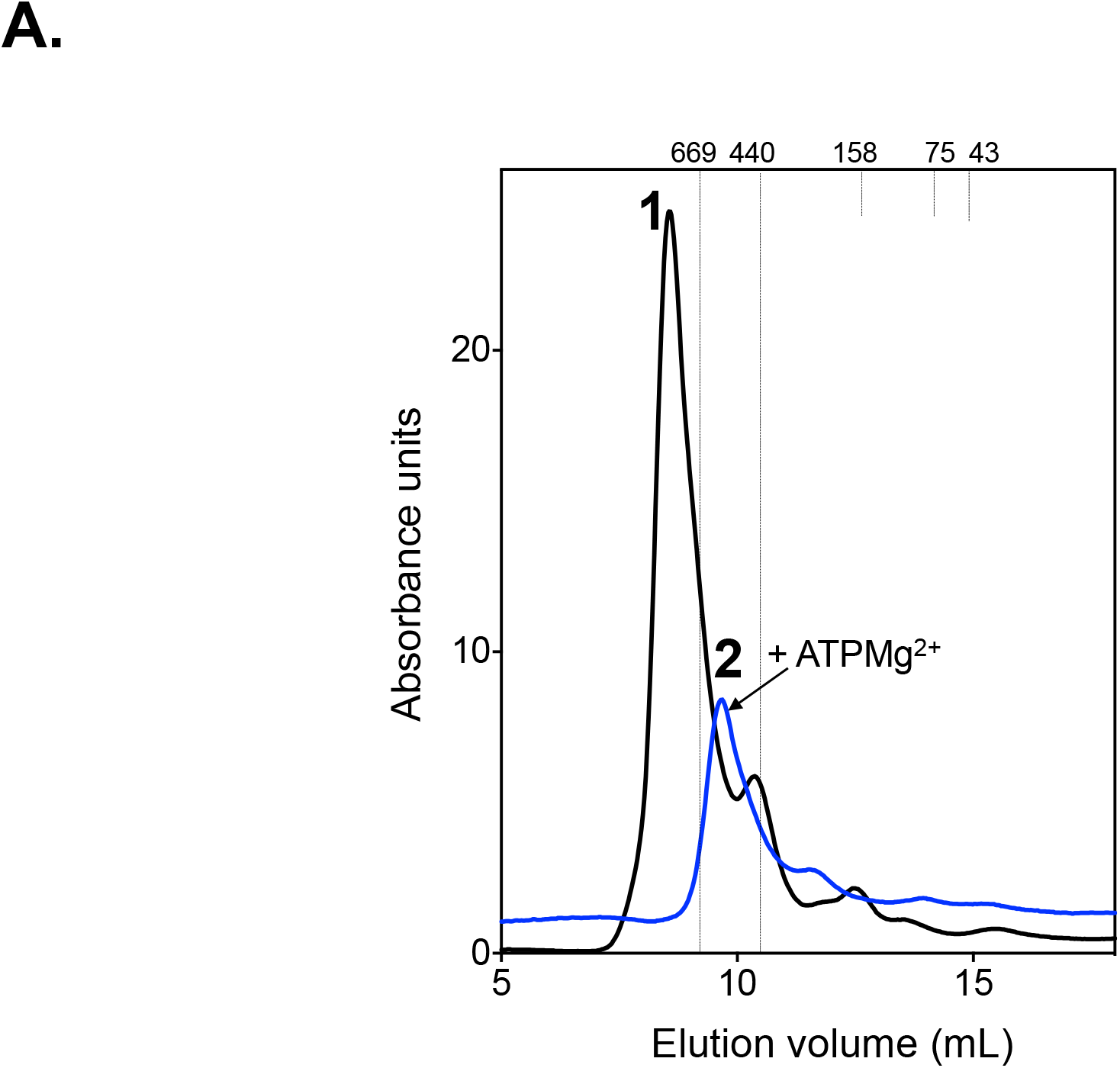
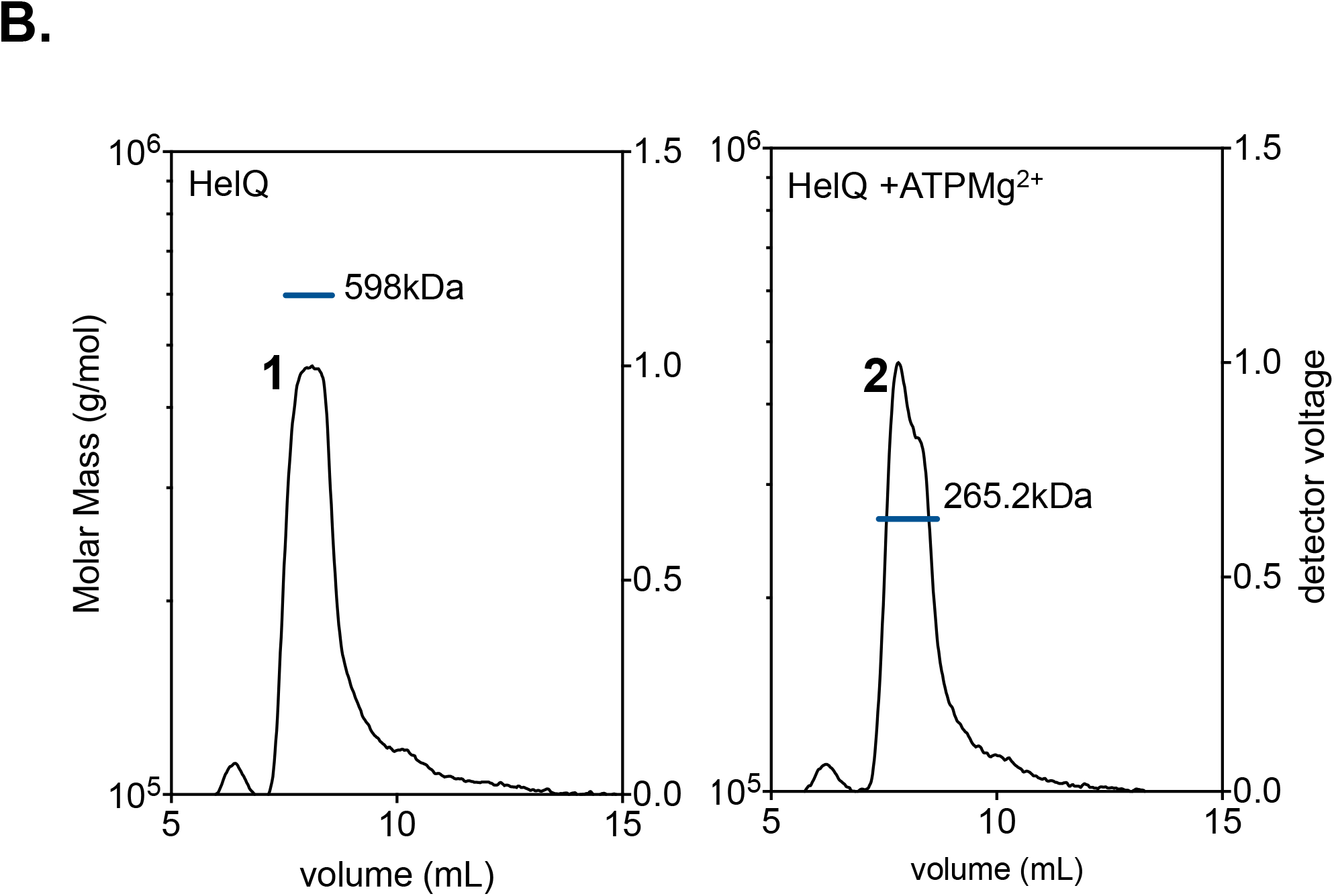
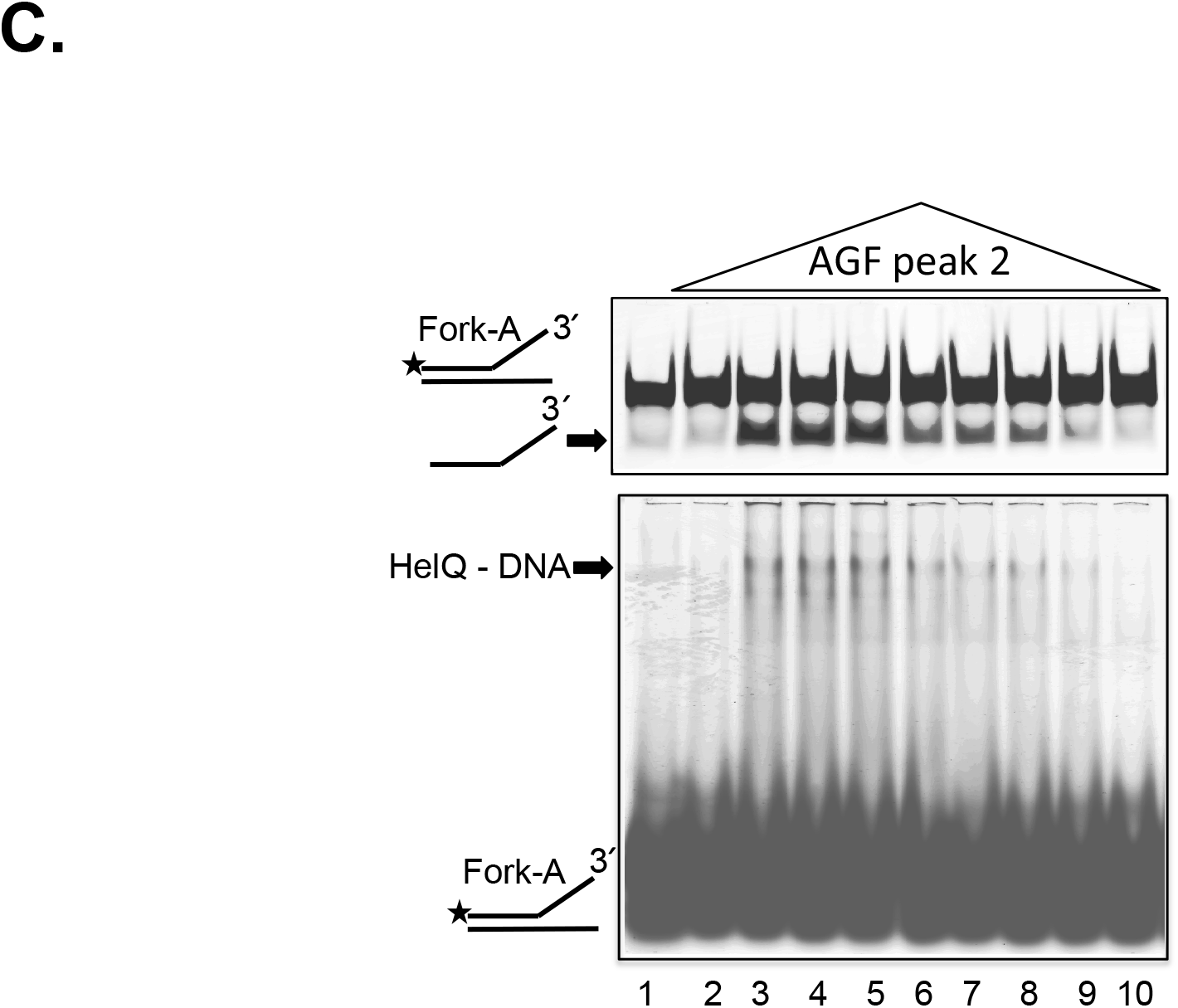
An active oligomeric state for HelQ. A. Combined traces for analytical gel filtration of pure human HelQ (200 μg) in standard buffer (black line, peak 1; Tris.HCl pH 8.0, 150 mM NaCl) compared to standard buffer supplemented by 5 mM ATP and 5 mM MgCl_2_ (blue line, peak 2). Molecular mass size markers were also run under both conditions giving the same elution volumes indicated by the dotted lines and numbers (KDa). B. SEC-MALS analysis of pure HelQ, giving molecular masses as indicated for peak 1 and peak 2 buffer conditions given in (A). Dynamic Light Scattering (DLS) measurements gave a mean poly-diversity index (PDi) of 0.28 for HelQ indicating acceptable monodispersal of HelQ particles. C. HelQ protein fractions collected from peak 2 buffer conditions (+ATP + Mg^2+^) were active as a DNA helicase when mixed with model fork-A DNA (25 nM, top gel panel helicase product indicated by an arrow), and this corresponded to fork-A binding by the same protein fractions in an EMSA, bottom gel panel. In the reactions, 20 μl of each 300 μl peak 2 fraction was used to assay for unwinding or binding to fork-A.

## DISCUSSION

We demonstrated that HelQ helicase and RPA heterotrimer form a stable ternary complex on DNA (Figure 2 and Supplementary Figures) and that RPA stimulates HelQ to unwind duplex DNA by displacing a high-affinity BamHI^E111A^ protein-DNA complex (K_d_ 2.95 × 10^−11^ M (42)) (Figures 2C and 2D and Supplementary). We present evidence that the N-terminus of HelQ (N-HelQ) interacts physically with RPA and that this interaction modulates RPA-DNA binding (Figure 3). We also discovered that 600 KDa HelQ multimers observed in previous work (26) are converted into dimers by ATP binding and that these bind to DNA and unwind it (Figure 4). ATP-dependent helicase activity of HelQ is activated when HelQ is bound to ssDNA, triggering ssDNA translocation with 3′ to 5′ directionality (26). In the assays presented here HelQ is provided with 3′-ended ssDNA to load onto fork-A or fork-B and catalyse unwinding through duplex DNA. Activation of HelQ helicase by ssDNA that is pre-bound with RPA would therefore require HelQ to gain access to ssDNA. Physical association of HelQ with RPA70 in cell extracts (23, 25), and as foci in replication stressed cells (32), provided evidence for recruitment of HelQ to RPA at sites of ssDNA. Our data show that this is also observable *in vitro* in the absence of DNA and when RPA is pre-bound to DNA, and leads to joint RPA-HelQ catalysed dissociation of a BamHI^E111A^ roadblock from duplex DNA, inferred from unwinding of Fork-B (Figure 2C). Significantly, in similar assays binding to RPA of a 417 amino acid protein fragment from the N-terminus of HelQ (N-HelQ) caused destabilised RPA-DNA binding (Figures 3D, 3E and 3F). This reveals a possible mechanism for how HelQ accesses ssDNA at RPA binding sites, triggering ATP-dependent translocation. In our EMSA assays eviction of RPA from DNA was observed as increasing amounts free-DNA substrate, corresponding to increased N-HelQ concentration, because N-HelQ is unable to bind to DNA (Figure 3E). However, binding of full length HelQ to ssDNA and RPA, preserving a HelQ-RPA-ssDNA ternary complex, would explain why free DNA was not observed in those EMSAs (e.g. in Figure 2A lane 7). Multiple DNA binding domains in human RPA heterotrimers confer dynamic DNA binding characteristics that are modulated during DNA repair and replication (44, 47). We propose that modulation of RPA-DNA binding induced by interaction with the HelQ N-terminal region reveals ssDNA for HelQ binding, leading to activation of HelQ helicase activity, summarised as a model in Figure 5. The amino acid sequence of the human N-HelQ region lacks homology to any other protein but is predicted be an intrinsically disordered protein, summarised in Supplementary Figure 7, joining a family of proteins that regulate protein-protein interactions in diverse aspects of cellular metabolism (48, 49). RPA may or may not remain bound to translocating HelQ, if so potentially providing motive power and/or processivity to displace other bound proteins during helicase unwinding of DNA. Further work, in particular utilizing HelQ and N-HelQ with fluorescent RPA proteins, will be needed to verify this model and determine mechanistic details.

**Figure 5.**
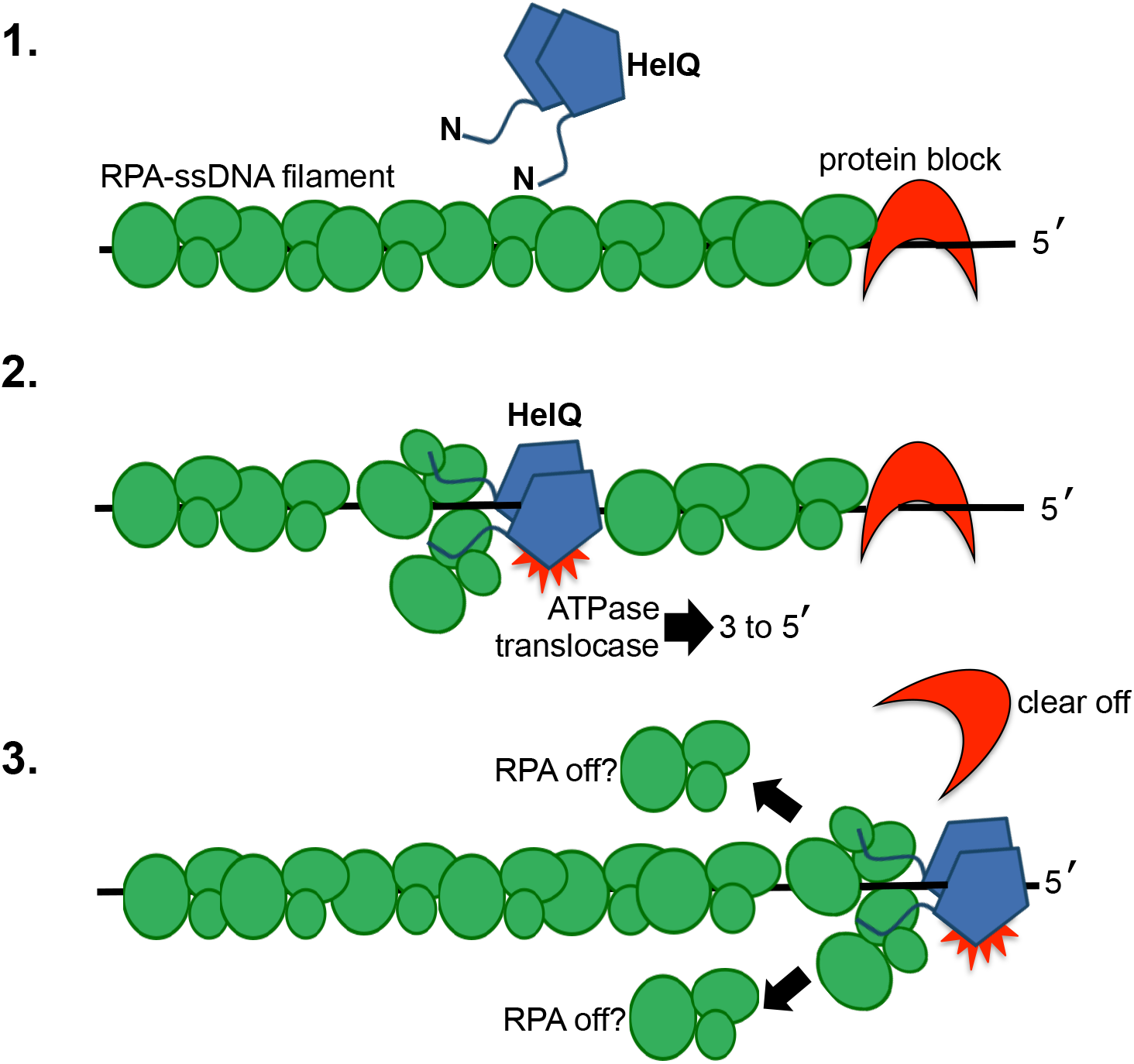
A model for HelQ-RPA interaction at ssDNA. 1. Aberrant ssDNA generated during replication stress is bound by RPA heterotrimers (green circles) and HelQ is present as a replication-coupled DNA repair helicase (blue pentagons). HelQ is represented as a dimer, based on SEC-MALS data (Figure 4) with the predicted intrinsically disordered N-terminal region indicated (Figure S7). In this scenario a protein-DNA complex (red crescent) would be a target for removal by HelQ. 2. Interaction between RPA and N-HelQ regions leads to modulated RPA-DNA binding that gives access for HelQ to load onto ssDNA, triggering ATPase activity and DNA translocation in the direction shown. 3. RPA-stimulated DA translocation by HelQ can be effective at displacing a protein-DNA complex. RPA may or may not remain dynamically associated with translocating HelQ.

RPA^Aro1^ did not stimulate HelQ to unwind DNA through BamHI^E111A^ (Figure 2C), but a HelQ-RPA^Aro1^-DNA ternary complex was observed, but with altered electrophoretic properties to HelQ-wild type RPA (Figure S3C). This indicates that helicase activity of HelQ-RPA requires precisely controlled DNA binding by RPA. RPA^Aro1^ mutations alter DNA binding by RPA70 that results in cellular defects in human cell DNA repair (50), raising the possibility that non-functioning HelQ-RPA complexes may be a direct contributory factor to repair phenotypes. *Helq* deletions cause genome instability and increased replication fork stalling in mammalian cells (24) (23, 25). Functional interaction of HelQ with RPA is therefore likely to be important for these processes, and adds to the list of RPA-interacting helicases and nucleases implicated in DNA repair (21, 51, 52). Notably, RPA stimulates the human Fanconi anemia pathway helicase FancJ through the same BamHI^E111A^ protein barrier as HelQ, but also through DNA strand damage or G4 DNA (42, 53, 54) which were not overcome by HelQ-RPA. *helq* and Fanconi anemia pathways are not epistatic and therefore HelQ-RPA and FancJ-RPA are likely to function in different repair contexts. Sensitivity of cells lacking *helq* to inter-and intra-strand DNA crosslinks (ICLs) implicates HelQ helicase activity in reactions that promote replication restart downstream of ICL unhooking reactions by nucleases. One requirement for HelQ may be analogous to a “fork-clearing role” described for bacterial UvrD, which removes proteins from sites of stalled replication in order to establish fork repair and fork reversal processes (55, 56). We observed using SEC-MALS that HelQ is helicase active as a protein of 265 kDa when bound to ATP-Mg^2+^, consistent with being a dimer. The N-HelQ fragment was monomeric and therefore dimerization must require interaction between HelQ helicase domains that comprise the remainder of the protein. The implications of dimerization to HelQ translocation mechanism and a role in replication-coupled repair will need detailed assessment using kinetics and biophysical techniques, but we note that helicase domains of DNA polymerase theta (PolQ), a close sequence homologue of HelQ, form a dimer of dimers (57) and that UvrD (58, 59) and RecQ form multimers in response to binding specific DNA structures (60).

## METHODS

### DNA substrates

DNA substrates are detailed in Supplementary Data Figure S6. Unmodified DNA oligonucleotides were purchased from SIGMA and were Cy5 labelled on 5’ ends as indicated in Figures. Chemically modified oligonucleotides were purchased from Eurofins (phosphorothioate DNA), EuroGenTech (Abasic DNA) or IDT (methylphosphonate DNA). Unless stated, enzymes for DNA manipulations were bought from New England Biolabs (NEB). Fork DNA substrates were made by annealing to room temperature overnight a 1.2:1 ratio of unlabelled: Cy5-labelled complementary oligonucleotides, after heating at 95°C for 10 minutes. DNA substrates were separated from unannealed oligonucleotides by gel electrophoresis through 10% w/v acrylamide Tris.HCl, borate and EDTA (TBE) gels, followed by excision of the desired substrate as a gel slice gel and soaking the slice overnight into Tris.HCl pH 7.5 + 150 mM NaCl to recover DNA. G4 Quadruplex was formed from a 50mer oligonucleotide following the method in (61), 3’ end-labelled using Aminoallyl-UTP-Cy5 (Jena Bioscience) incubated with TdT.

### Proteins

HelQ and the N-terminal HelQ fragment (N-HelQ) were over expressed using the Bac-to-Bac baculovirus expression system from Gibco BRL in SF9 insect cells, each with N-terminal (His)_6_, SUMO and Strep-Flag tag (25). SF9 cells were cultured in LONZA Insect Xpress protein-free culture media with L-glutamine supplemented with pluronic acid, penicillin-streptomycin solution and amphotericin B solution. HelQ protein over-expression was optimised at 5 μl virus per 1×10^6^ SF9 cells for 48 hours at 27°C. The N-terminal region of HelQ was produced by over-expression of HelQ for prolonged periods, typically 72 hours at 27°C, resulting in natural protein degradation and the formation of a stable tagged HelQ fragment, summarised in Supplementary Figure S5. This fragment (N-HelQ) comprised HelQ amino acids 1 – 417 as verified by mass spectrometry.

For purification of full-length HelQ all stages used columns and buffers refrigerated to 4°C with Coomassie staining of SDS-PAGE gels for identification of HelQ containing fractions. Biomass was resuspended in lysis buffer (150 mM Tris pH8, 150 mM NaCl, 20 mM imidazole, 10% glycerol) containing COMplete EDTA free protease inhibitor tablets and stored at −80°C until purification. Cells were thawed on ice over several hours before lysis on ice by sonication at 80% pulsed for 1 min per 5 ml biomass. Subsequent soluble proteins were fractionated by precipitation with 0 – 50 % solid ammonium sulphate added on ice with gentle agitation, at a rate of 12 g in 40 minutes. The resulting protein precipitate was recovered by centrifugation and resuspended in 50mM NaCl, 20mM Imidazole, 50mM Tris pH8, 10% glycerol, 1mM TCEP to dissolve precipitated proteins. Solubilised proteins were loaded onto a 5 ml NiCl_2_ charged Ni-NTA column that was washed with Ni-NTA-A buffer (20 mM Tris.HCl pH 8.0, 10% w/v glycerol, 1 M NaCl, 10 mM imidazole, COMplete EDTA free protease inhibitor tablet). (His)_6_HelQ protein that eluted in a gradient of imidazole at approximately 100 – 200 mM imidazole 10 - 20% was pooled protein and dialysed over-night at 4°C into 150 mM NaCl, 10% w/v glycerol, 50 mM Tris.HCl pH 8.0. This was loaded onto a 5ml heparin column, washed in HepA buffer (50 mM Tris.HCl pH 8.0, 10% w/v glycerol, 150 mM NaCl, 1 mM TCEP, COMplete EDTA free protease inhibitor tablet) and bound HelQ eluted from heparin between 100 – 400 mM NaCl. HelQ fractions were diluted with cold buffer HepA to reduce NaCl to approximately 200 mM and the protein was next loaded onto a 1ml anion exchange Q-sepharose column. This column was washed in buffer HepA and HelQ eluted in a gradient of increasing NaCl concentration at approximately 350 – 600 mM NaCl. Pooled HelQ protein was dialysed for 3 hours at 4°C into 30% w/v glycerol, 5 mM DTT, 150 mM NaCl, 50 mM Tris.HCl pH 8.0 and aliquotted for storage at −20°C. Protein concentration was calculated using Bradford’s reagent. N-HelQ purification was carried out as described above except that the heparin column step was not included. Replication Protein A (RPA) was purified at the Research complex in Harwell following previously described methods (62). Marc Wold (University of Iowa) kindly provided purified RPAAro1 protein.

### Standard Methods for Protein-DNA assays

Electrophoretic Mobility Shift Assays (EMSAs) were in buffer HB (100 mM Tris pH 7.5, 500 μg/ml BSA and 30% glycerol) containing fresh dithiothreitol (DTT, 25 mM). Protein-DNA reactions were incubated at 37°C for 10 minutes in 20 μl reaction volumes. Protein dilutions were made resulting in final concentrations described in the results, adding a final volume of 2 μl of protein to reactions. Reactions were loaded directly onto 5% w/v acrylamide TBE gels for electrophoresis at 150 volts for two hours in Protean II tanks in TBE buffer. In EMSA reactions containing RPA and HelQ, RPA was pre-incubated with DNA for 2 min. at room temperature prior to adding HelQ.

Helicase unwinding assays were in the buffer HB buffer supplemented with 5mM magnesium chloride and 5 mM ATP. Cy5 labelled DNA substrate (25 nM) was used with addition of ‘cold-trap’ unlabelled oligonucleotide to 2.5 nM. Helicase reactions were stopped by addition of buffer STOP comprising 2.5 % w/v SDS, 200 μM EDTA and 2 μg/μl of proteinase K. Reaction products were assessed after electrophoresis through 10% acrylamide TBE gels. Gels were imaged using FLA-3000 or Typhoon machine and quantified using ImageJ, GelEval and Prism software.

### BamHI-EIIIA Roadblock Assays

BamHI-EIIIA roadblock experiments were conducted as above with pre-incubation of the substrate with BamHI-EIIIA at described concentrations at room temperature for 15 min. prior to addition of reaction components and HelQ protein. Experiments including RPA were carried out with an additional incubation step prior to addition of HelQ with RPA at described concentrations at 37°C for 10 min.

### Analysis of N-HelQ in structural databases

The amino acid sequence of N-HelQ was confirmed by mass spectrometry and the tertiary structure of this region was predicted with PHYRE2 (63). Resulting predicted structures were analysed and superimposed with DALI (64) giving a structural homology match for Brr2 PWI from PDB accession 5DCA.

### RPA-HelQ protein pull-down assays

Protein pull down assays between HelQ and RPA were carried out using streptactin resin to trap (His)_6_-strep-tagged HelQ in a gravity flow streptactin-sepharose column (IBA solutions). HelQ and RPA were pre-incubated in a 1:1 molar ratio on ice for 30 minutes in buffer comprising 50mM Tris pH7.5, 10% glycerol, 25mM DTT, 150mM NaCl and 0.02% w/v Tween 20. Proteins were applied to streptactin columns equilibrated in the same buffer at 4°C, with flow-through samples being collected and re-applied to the column three times for maximal HelQ binding. Columns were then washed with one column volume of equilibration buffer prior to elution of bound HelQ using the same buffer containing 2.5 mM desthiobiotin. Collected samples were analysed by SDS-PAGE 8% w/v acrylamide gels.

### Analysis of HelQ by size exclusion chromatography and SEC-MALS

Size exclusion chromatography-multi-angle static light scattering (SEC-MALS) was carried out using a Wyatt Dawn 8+ 1260 Infinity II series in a system equipped with a Superdex-200 column size exclusion column. Pure HelQ protein (200 mL of 2.4 mM, calculated as monomer concentration) was applied to the column at a flow rate of 1ml/min, after pre-incubated of HelQ in buffer for 10 minutes at 37°C. Protein mass was calculated using Wyatt DAWN^®^ HELEOS^®^ II MALS, using a dn/dc of 0.185. The resulting chromatograms were analyzed using ASTRA^®^ software, V.6.1.2.84 (Wyatt Tech Corp). The analyses used three conditions, as described in results. Standard buffer comprised 50mM Tris-HCl pH 8.0, 150mM NaCl and 10% glycerol. For + ATP-Mg^2+^ reactions, standard buffer was supplemented with 0.2mM ATP and 0.2mM MgCl_2_ and HelQ was pre-incubated in 5mM MgCl_2_ and 5mM ATP at 37°C for 10 minutes. For reactions + ssDNA, HelQ was pre-incubated in standard buffer + ATP-Mg^2+^ and with the addition of 5 μM of MW14 ssDNA, details of which are in Supplementary Figure S6.

### ATPase assays

Measurement of HelQ ATPase activity used the malachite green dye assay. Reactions in 20 uL volumes contained protein mixed with magnesium and ATP (5 mM each) and either no DNA, MW14 oligonucleotide (25nM) ssDNA, or M13 single stranded circular DNA (200 ng), as stated in results and Supplementary figures. Reactions were at 37°C for 10 minutes, concluded by addition of 80 μL of detergent-free water and 800 μL of dye reagent for 2 minutes at room temperature, followed by addition of a 34% aqueous solution of sodium citrate to allow colour development for a further 30 minutes. Concentration of liberated phosphate was measured by absorbance at 660 nm against a no protein blank and a calibration curve of known phosphate concentrations. Dye reagent was a 3:1 ratio of stock solutions of malachite green: ammonium molybdate. Malachite green stock was 0.045% (weight to volume) malachite green in water, and ammonium molybdate to 4.2% (weight to volume) in 4M HCl.

## ACKNOWLEDGEMENTS

We are grateful to the following for kind donations of proteins: Marc Wold and Robert Brosh Jr. for RPA^ARO1^, Peter McGlynn for *E. coli* SSB and New England Biolabs for BamHI^E111A^. We thank the Oxford Protein Production Facility (OPPF) at Harwell for initial help generating recombinant human proteins. The work was supported by PhD studentships from The University of Nottingham (SJN) and The BBSRC DTP scheme (TJ).

