## Supplementary Data Figures and Legends for "Interaction of HelQ helicase with RPA modulates RPA-DNA binding and stimulates HelQ to unwind DNA through a protein roadblock"

#### Supplementary Figure Legends

**Figure S1. Purified human His-strep tagged HelQ is a ssDNA dependent ATPase and helicase.**

A. His-Strep-HelQ purified in this work, with expected molecular mass of 141 KDa, imaged from a Coomassie stained SDS-PAGE gel.
B. ATPase activity of purified HelQ protein from (A) (100 nM) mixed with M13 ssDNA (250 ng) or in the absence of DNA as indicated.
C. Helicase activity of purified HelQ protein from (A) (100 nM) mixed with fork-A DNA (see Figure S6, 25 nM) in the presence or absence of 5 mM magnesium chloride and 5 mM ATP, as indicated. The star on fork-A shows the site of Cy5 labelling of one DNA strand.

**Figure S2.**

A. Graph showing unwinding of DNA fork-A and its chemically modified derivatives as a function of HelQ protein concentration. The forks used (each 25 nM) are the same as in Results Figure 1 and HelQ was added at 5, 10, 20, 40, 80 and 160 nM. The graphs represent mean unwinding of two independent experiments for each fork, with error bars showing standard deviation from the mean.
B. Summary EMSAs of HelQ binding to each fork-A from (A)(25 nM) using HelQ at the same concentrations.
C. EMSA summarising binding of BamHI$^{E111A}$ protein at 5, 10, 20, 40, 80 and 160 nM to fork-B DNA (25 nM).

**Figure S3.**

A. EMSA summarising binding of HelQ (50 and 100 nM), RPA (10 and 25 nM) and RPA-HelQ (100-25nM) to fork-A DNA (25 nM) in lanes 1-6. *E. coli* SSB (5 and 15 nM) also binds to fork-A DNA (lanes 7 and 8) and addition of HelQ (100 nM) gives a HelQ complex and an SSB complex but no observable ternary complex (lane 9).

B. Graph showing helicase unwinding of fork-B DNA in various conditions, as labelled, using HelQ at 5, 10, 20, 40, 80 and 160 nM. HelQ unwinds fork-B in the absence of BamHI$^{E111A}$ ("HelQ unwinding") but pre-incubation of fork-B with BamHI$^{E111A}$ (160 nM) reduces this activity ("+BamHI"). Addition of RPA (5 nM), but not SSB (7.5 nM) or RPA$^{Aro1}$ (5 nM), restores HelQ catalysed unwinding of fork-B bound by BamHI$^{E111A}$.

C. EMSA summarising different ternary complex formation by RPA-HelQ (lanes 5 and 6, arrow 1) compared to RPA$^{Aro1}$-HelQ (lanes 7 and 8, arrow 2). HelQ binding alone is shown in lanes 3 and 4. Protein concentrations are the same as in (A). The image shown is taken from a single gel, with the relevant lanes tiled together to remove intervening unrelated gel lanes.

D. Summary HelQ helicase unwinding of fork-B pre-bound by BamHI$^{E111A}$ in the presence of ATP-$Mg^{2+}$ (top panel) and in the absence of ATP-$Mg^{2+}$ (bottom panel). In each case HelQ was added to 10, 20, 40, 80 or 160 nM and RPA at 5 nM. In the bottom panel lane 9 indicates addition of only RPA to fork-B BamHI$^{E111A}$. Lanes labelled B are boiled fork-B DNA.

E. HelQ (160 nM) ATPase activity measured in the presence of RPA (5 nM) or its absence. In all reactions MW14 ssDNA was added to 25 nM to stimulate HelQ ATPase activity.

F. Summary unwinding of G4 Quadruplex DNA (25 nM, see Figure S6) by human HelQ (5, 10, 20, 40, 80 and 160 nM) compared to purified *E. coli* RecQ (same concentrations), in the top panel. The bottom panel shows weak HelQ helicase unwinding of the same substrate in the absence of RPA and when RPA (5 nM) was pre-incubated with DNA.

G. Summary helicase unwinding of Fork-AP DNA (25nM, Fork-A with a single abasic site modification, Figure S6) in the presence of RPA (5 nM) or with no RPA. HelQ was used at 40, 80 and 160 nM in these reactions.

**Figure S4**

A. Cartoon representation of the *S. cerevisiae* Brr2 protein structure (PDB code 5DCA) highlighting the non-canonical PWI-like fold identified in the N-terminal region of this protein, superimposed with a model for HelQ as in results Figure 3. The highlighted region is shown as a structural alignment between human Brr2 (Hsa), *S. cerevisiae* Brr2 (Sce) and the amino acid sequence 240 – 348 at the N-terminus of HelQ (N-HelQ). Boxed is a single tract of high sequence conservation between the three proteins that is thought to be important for PWI function.

B. The top panel is a coomassie stained SDS-PAGE gel showing a sample of spontaneously degrading HelQ protein during purification, with N-HelQ fragment highlighted with *, and purified N-HelQ fragment derived from degradation of full length HelQ. The bottom panels summarise that N-HelQ (10, 100, 500 nM) did not bind to fork-A DNA (25 nM) in EMSAs and does not unwind fork-A DNA (25 nM), compared to active full length HelQ used at 10, 50 and 150 nM in these assays.

C. EMSAs summarising binding of RPA and N-HelQ to fork-A DNA (25 nM) in EDTA and in 5 mM magnesium chloride, or to ssDNA oligonucleotide in standard EMSA assay conditions, as indicated. RPA was used at 15 nM and N-HelQ at 200 nM.

**Figure S5**

A. SEC-MALs trace from HelQ in buffer containing $MgCl_2$ and ATP after pre-binding HelQ with MW14 ssDNA (25 nM).

**Figure S6**

Summary of DNA substrates used in this work. In part A and part B, Fork-A and Fork-B are shown as annealed oligonucleotides. In fork-B the BamHI$^{E111A}$ cognate sequence is in bold. In part C the table gives DNA sequences of each oligonucleotide used to generate the substrates described. In bold underlined indicates the location of the phosphodiester bond chemical modifications in Fork-Me and Fork-S and in Fork-AP the cytosine nitrogenous base was removed where indicated.

**Figure S7**

Interrogation of human HelQ amino acid sequence through databases for prediction of intrinsically disordered proteins highlighted prediction for >75% likelihood that N-HelQ sequences shown in bold are intrinsically disordered. This analysis was carried out using http://d2p2.pro (1).

1. Oates ME, Romero P, Ishida T, Ghalwash M, Mizianty MJ, et al. 2013. *Nucleic Acids Res* 41: D508-16

Figure S1

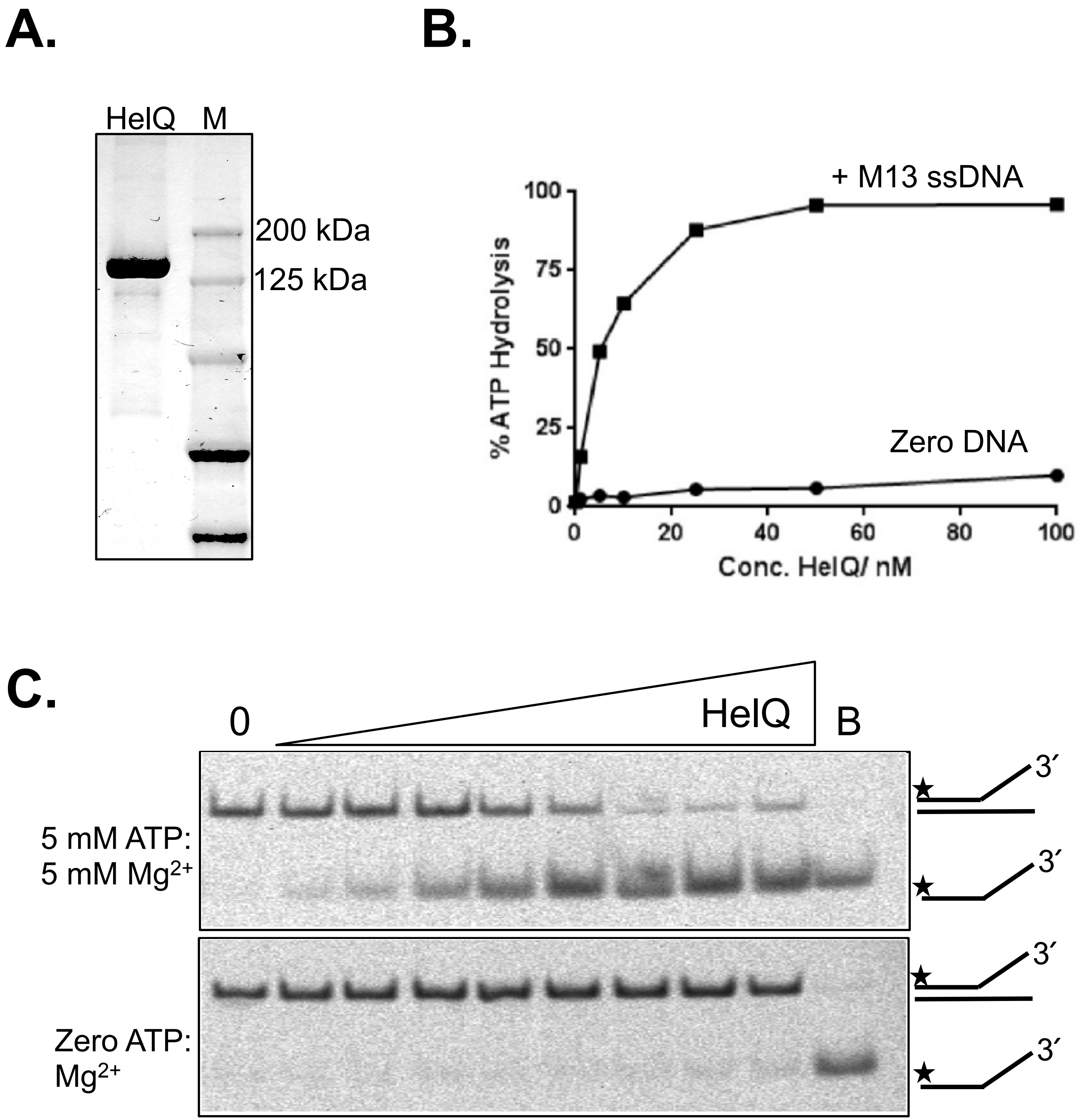
A.
HelQ
M
200 kDa
125 kDa
B.
+ M13 ssDNA
Zero DNA
% ATP Hydrolysis
100
75
50
25
0
0
20
40
60
80
100
Conc. HelQ/ nM
C.
0
HelQ
B
5 mM ATP:
5 mM Mg2+
Zero ATP:
Mg2+
3′
3′
3′
3′

Figure S2

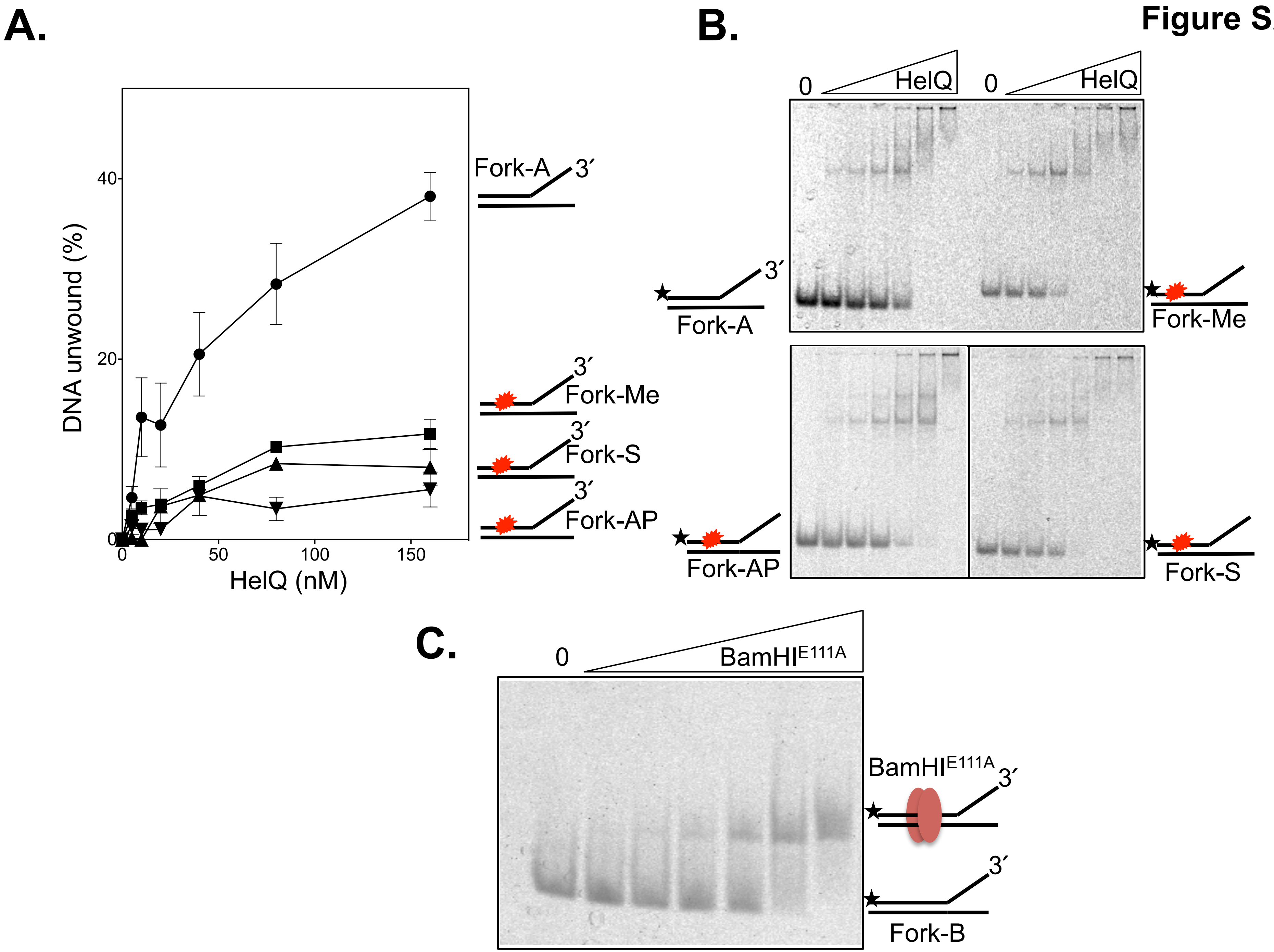
A.
DNA unwound (%)
HelQ (nM)
0
20
40
0
50
100
150
Fork-A
3′
Fork-Me
Fork-S
Fork-AP
B.
0
HelQ
Fork-A
Fork-Me
Fork-AP
Fork-S
C.
0
BamHI$^{E111A}$
3′
Fork-B

Figure S3

A.

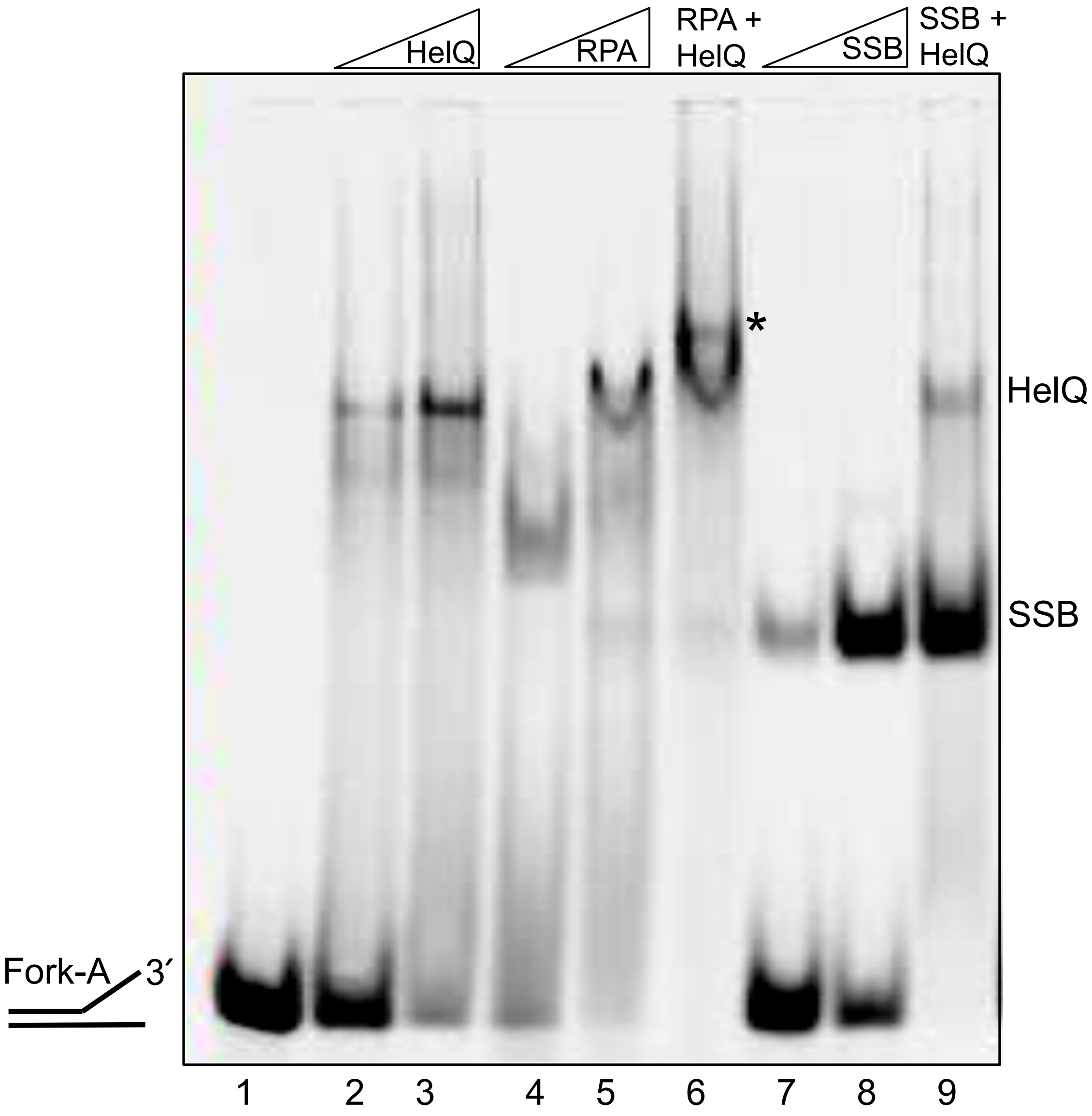

**B.**

**Figure S3**

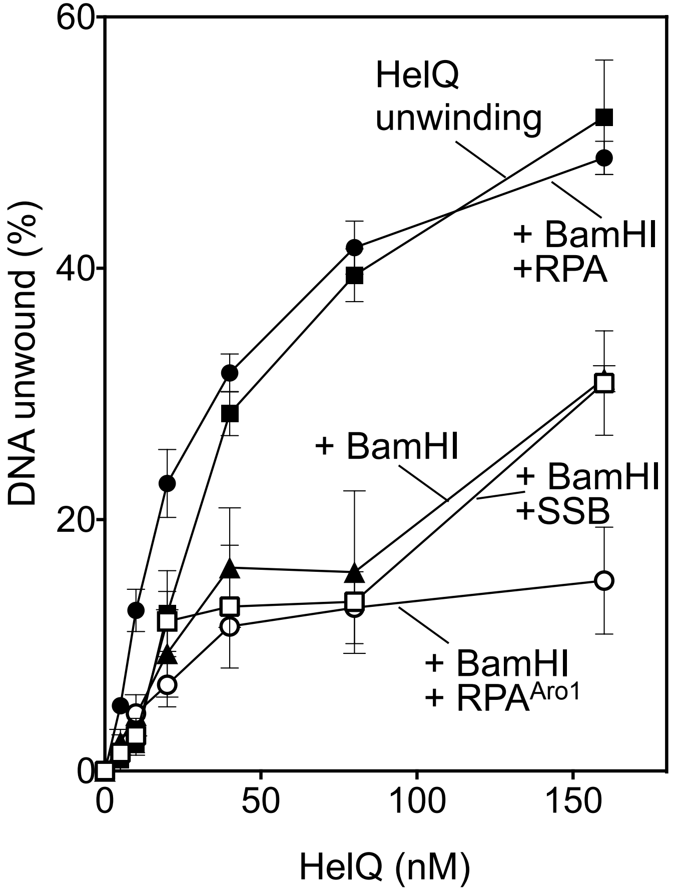

DNA unwound (%)
60
40
20
0
0
50
100
150
HelQ (nM)
HelQ
unwinding
+ BamHI
+RPA
+ BamHI
+ BamHI
+SSB
+ BamHI
+ RPA^Aro1

**C.**

**Figure S3**

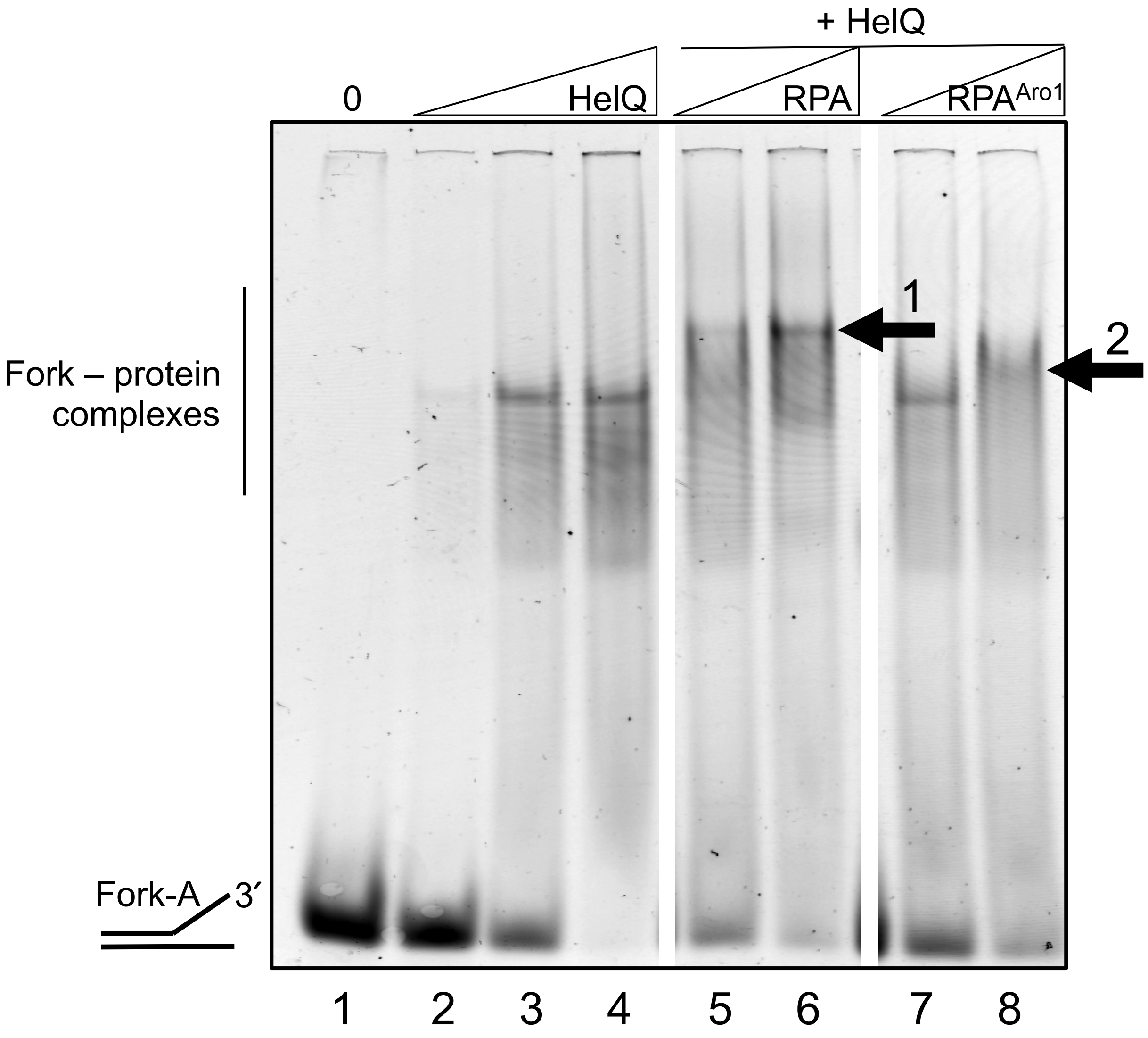
+ HelQ
0
HelQ
RPA
RPA^Aro1
Fork – protein complexes
1
2
Fork-A
3′
1 2 3 4 5 6 7 8

Figure S3

D.

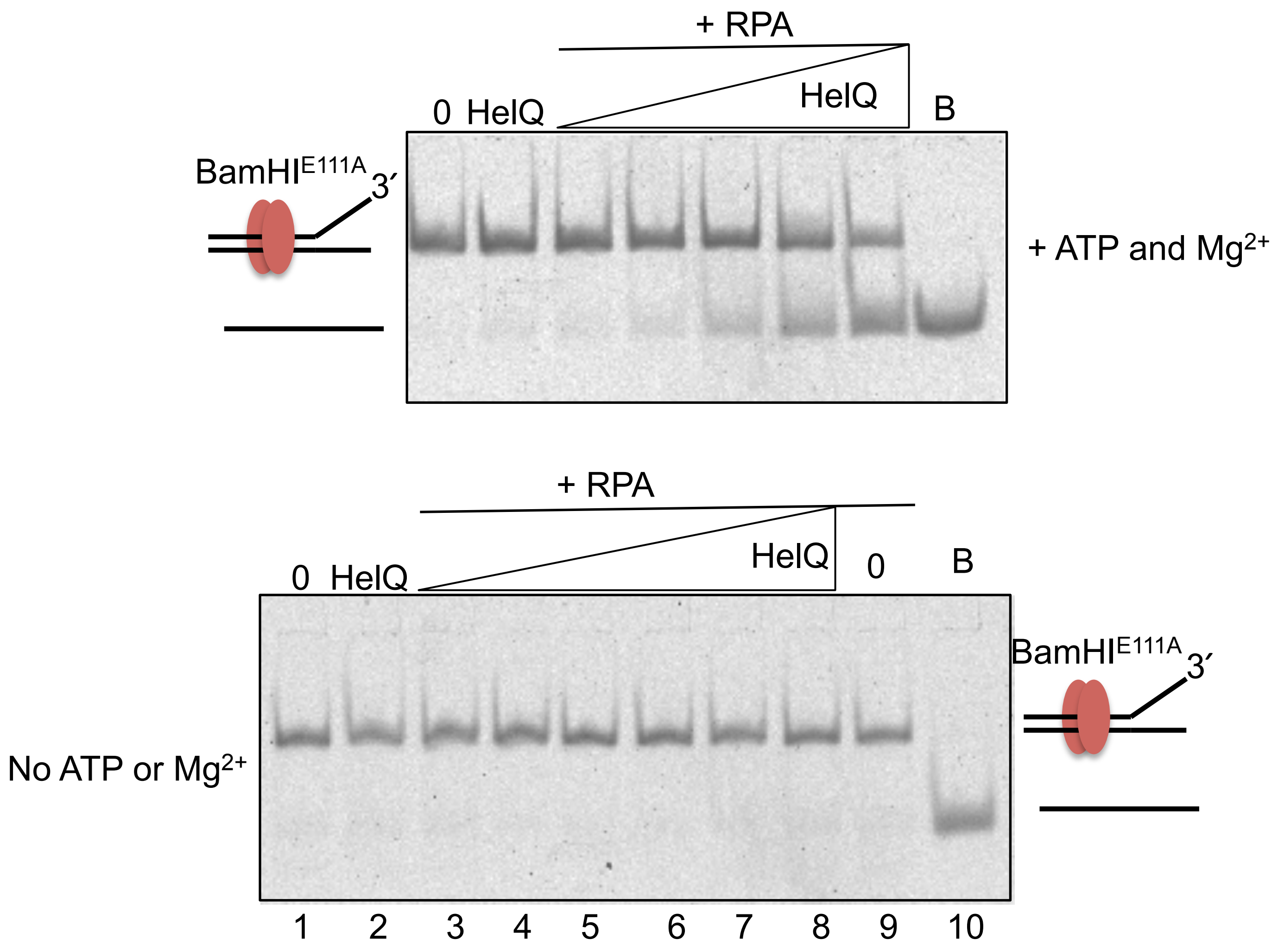
+ RPA
0 HelQ
HelQ
B
BamHI$^{E111A}$
3′
+ ATP and Mg$^{2+}$
+ RPA
0 HelQ
HelQ
0
B
BamHI$^{E111A}$
3′
No ATP or Mg$^{2+}$
1 2 3 4 5 6 7 8 9 10

# E.

**Figure S3**

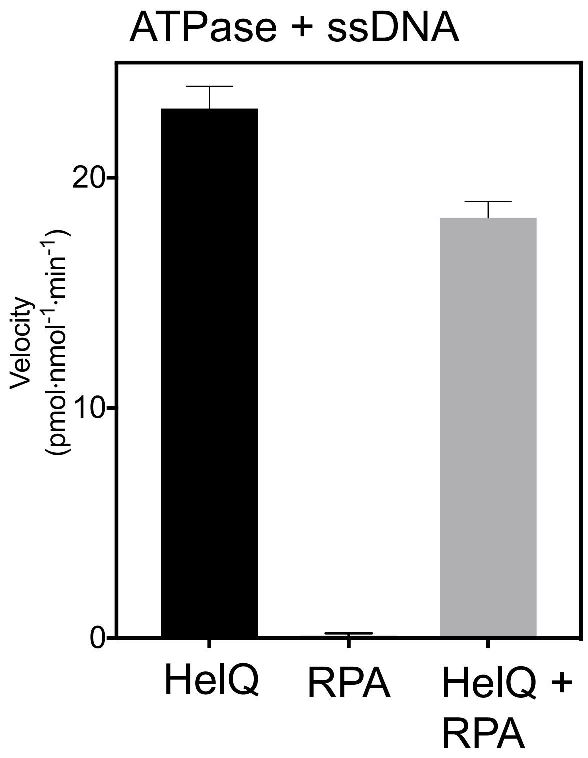

ATPase + ssDNA
Velocity
(pmol·nmol-1·min-1)
20
10
0
HelQ
RPA
HelQ +
RPA

**F.**

**Figure S3**

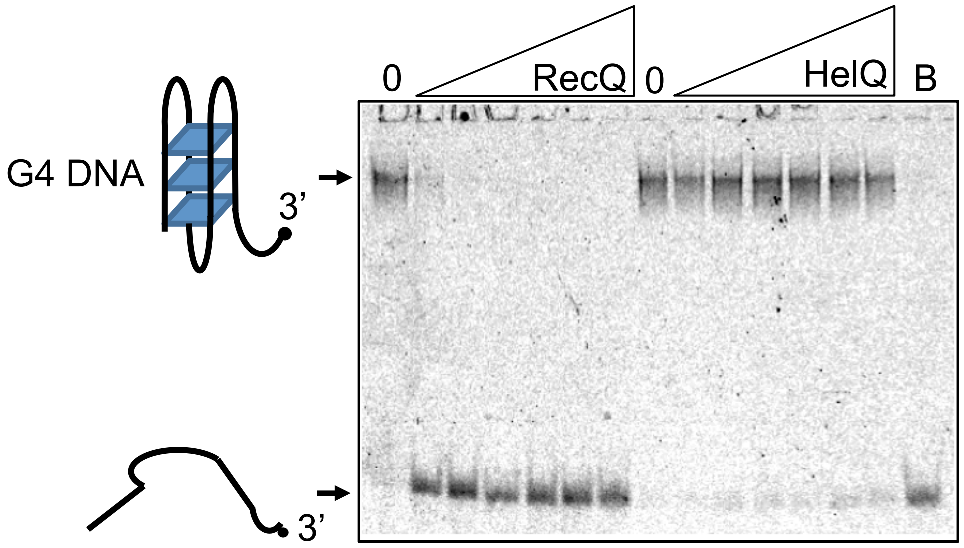
0
RecQ
0
HelQ
B
G4 DNA
3'
3'

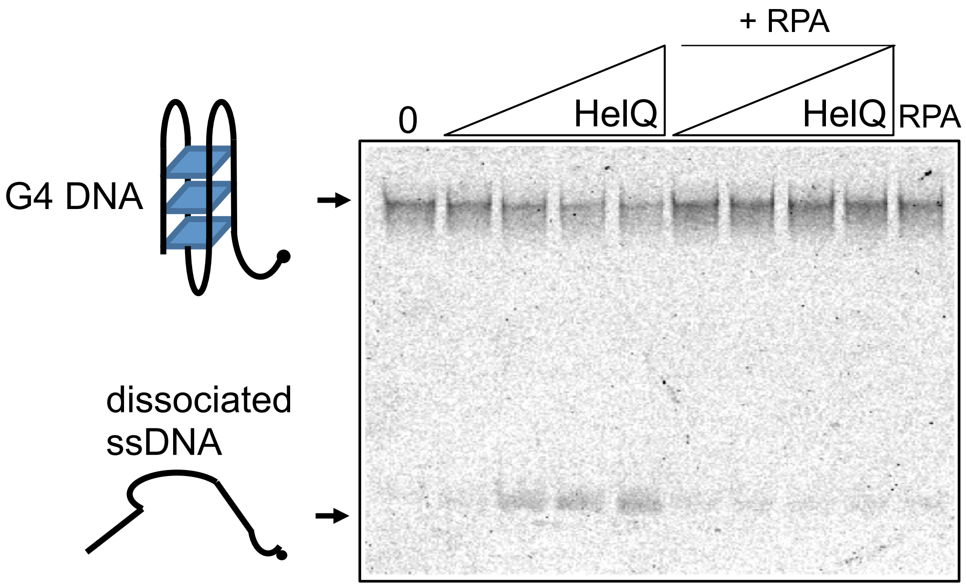
+ RPA
0
HelQ
HelQ
RPA
G4 DNA
dissociated
ssDNA

G.

Figure S3

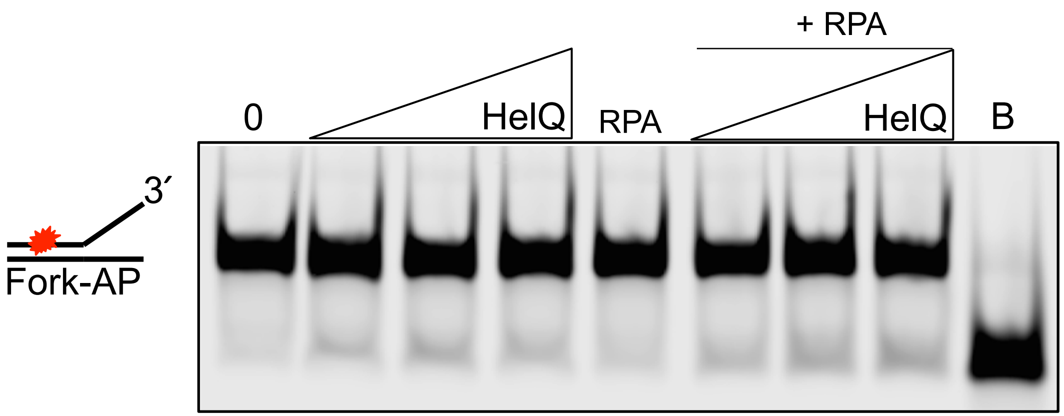

### Figure S4

**A.**

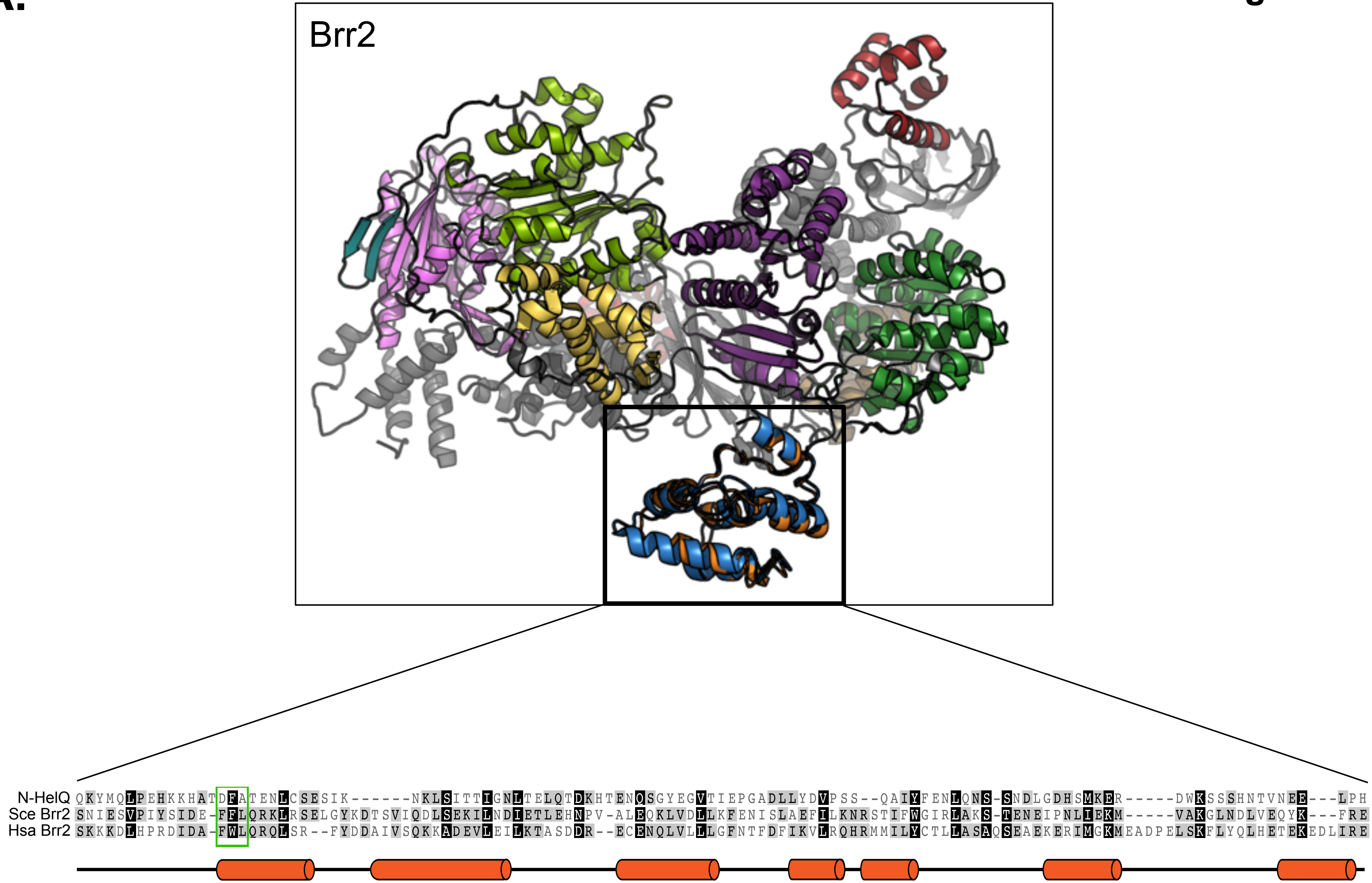

Brr2
N-HelQ QKYMQLPEHKKHATDFATENLCSESIK------NKLSITTIGNLTELQTDKHTENQSGYEGVTIEPGADLLYDVPSS--QAIYFENLQNS-SNDLGDHSMKER-----DWKSSSHNTVNEE---LPH
Sce Brr2 SNIESVPIYSIDE-FFLQRKLRSELGYKDTSVIQDLSEKILNDIETLEHNPV-ALEQKLVDLLKFENISLAEFILKNRSTIFWGIRLAKS-TENEIPNLIEKM-----VAKGLNDLVEQYK---FRE
Hsa Brr2 SKKKDLHPRDIDA-FWLQRQLSR--FYDDAIVSQKKADEVLEILKTASDDR--ECENQLVLLLGFNTFDFIKVLRQHRMMILYCTLLASAQSEAEKERIMGKMEADPELSKFLYQLHETEKEDLIRE

### Figure S4

**B.**

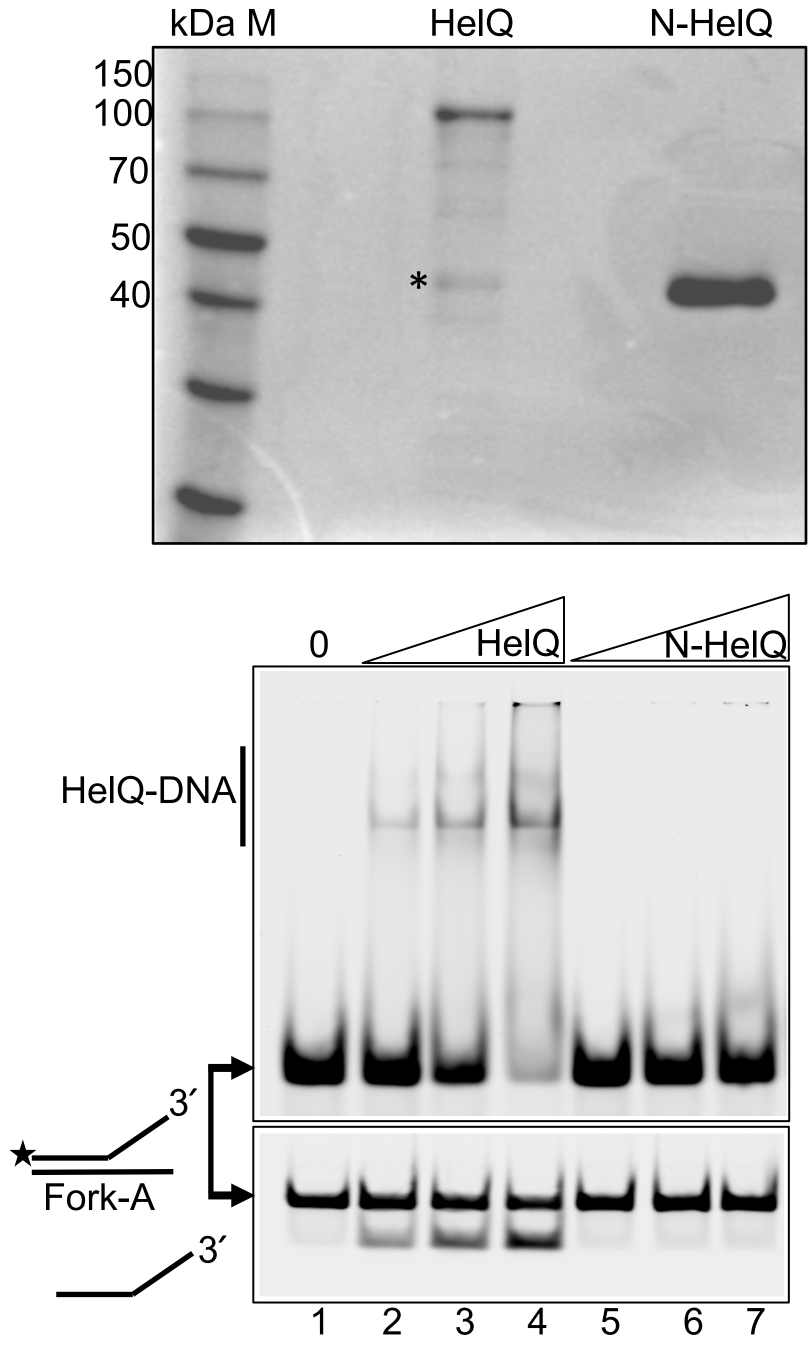
kDa M
HelQ
N-HelQ
150
100
70
50
40
*
0
HelQ
N-HelQ
HelQ-DNA
3′
Fork-A
3′
1
2
3
4
5
6
7

**C.**

**Figure S4**

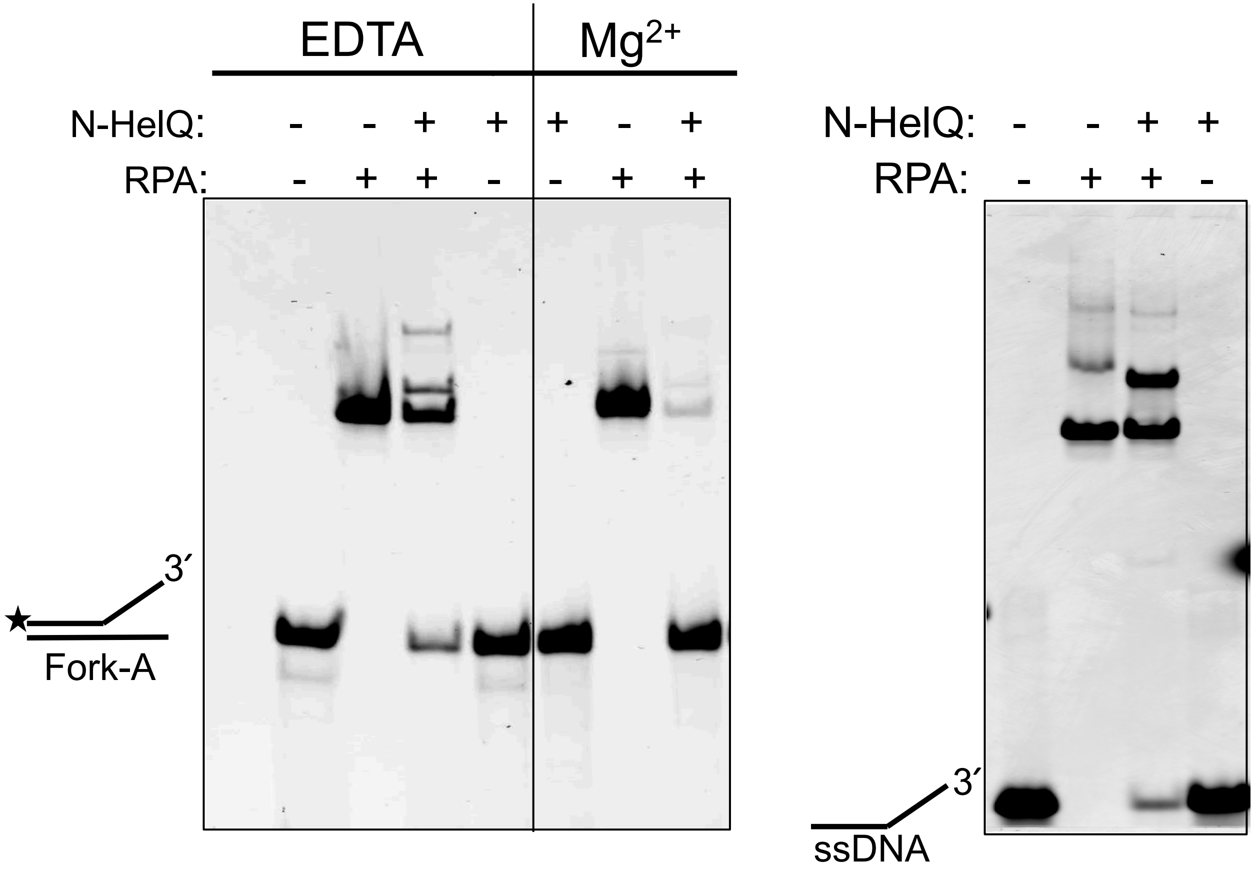

**Figure S5**

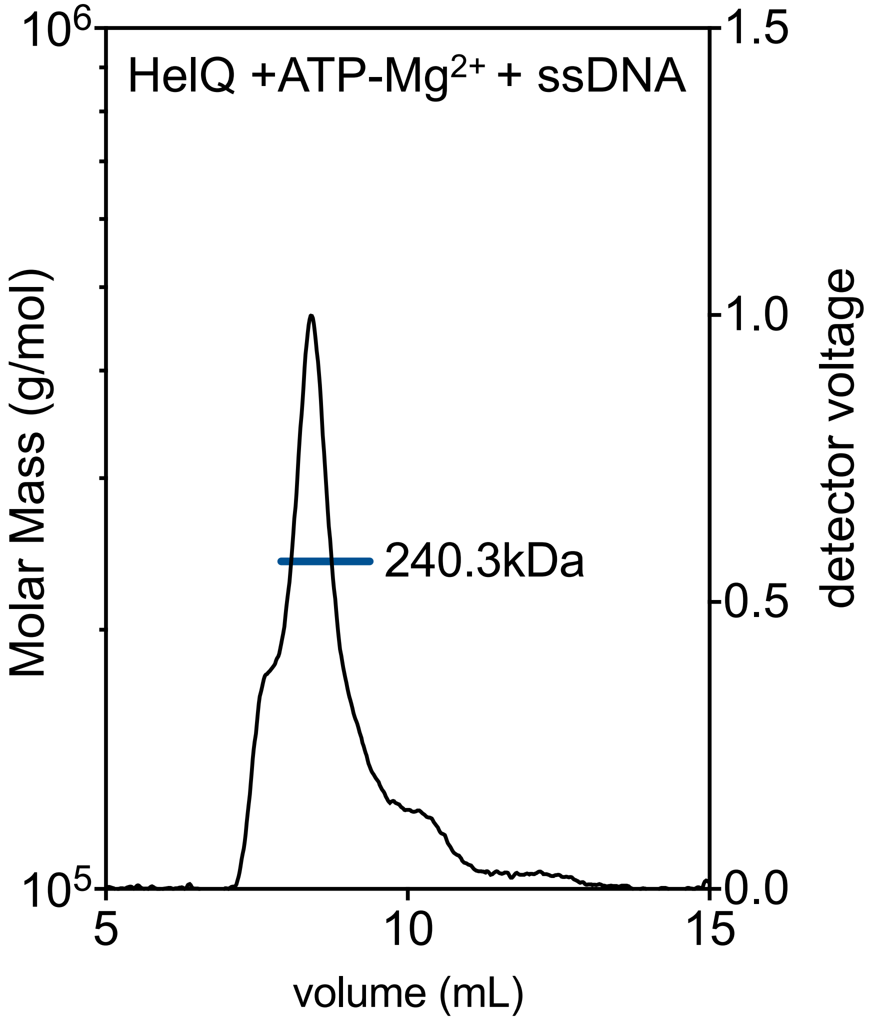
HelQ +ATP-Mg2+ + ssDNA
240.3kDa
Molar Mass (g/mol)
10^6
10^5
detector voltage
1.5
1.0
0.5
0.0
5
10
15
volume (mL)

# A.

**Figure S6**

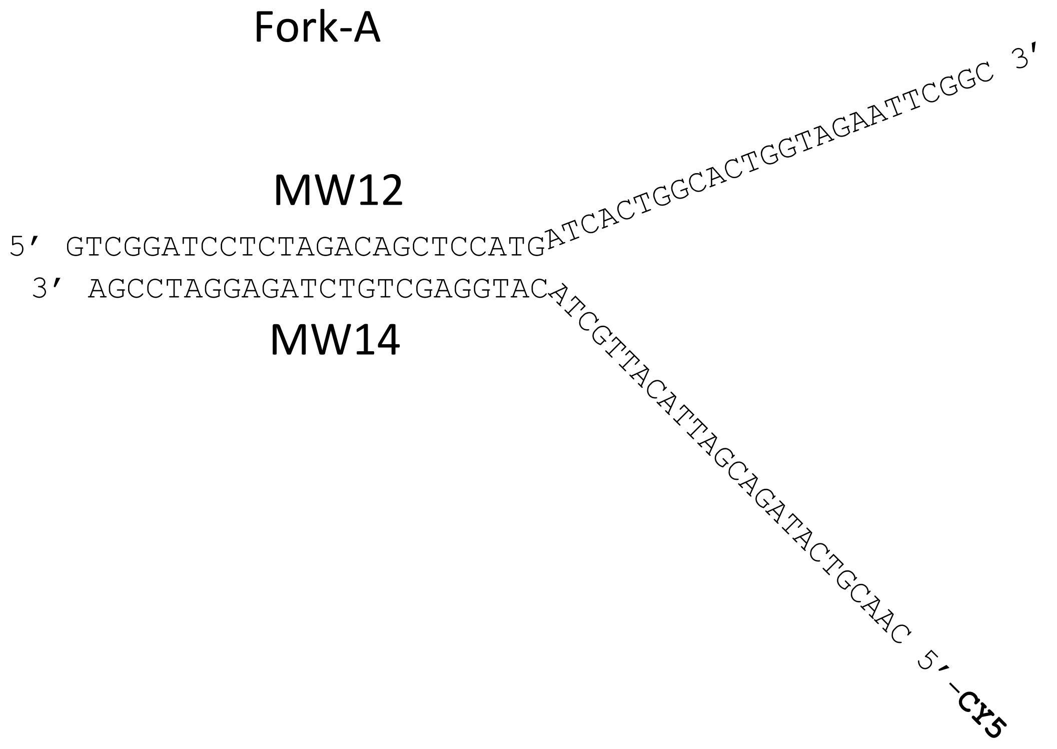
Fork-A
MW12
5′ GTCGGATCCTCTAGACAGCTCCATGATCACTGGCACTGGTAGAATTCGGC 3′
3′ AGCCTAGGAGATCTGTCGAGGTACATCGTTACATTAGCAGATACTGCAAC 5′-CY5
MW14

**B.**

**Figure S6**

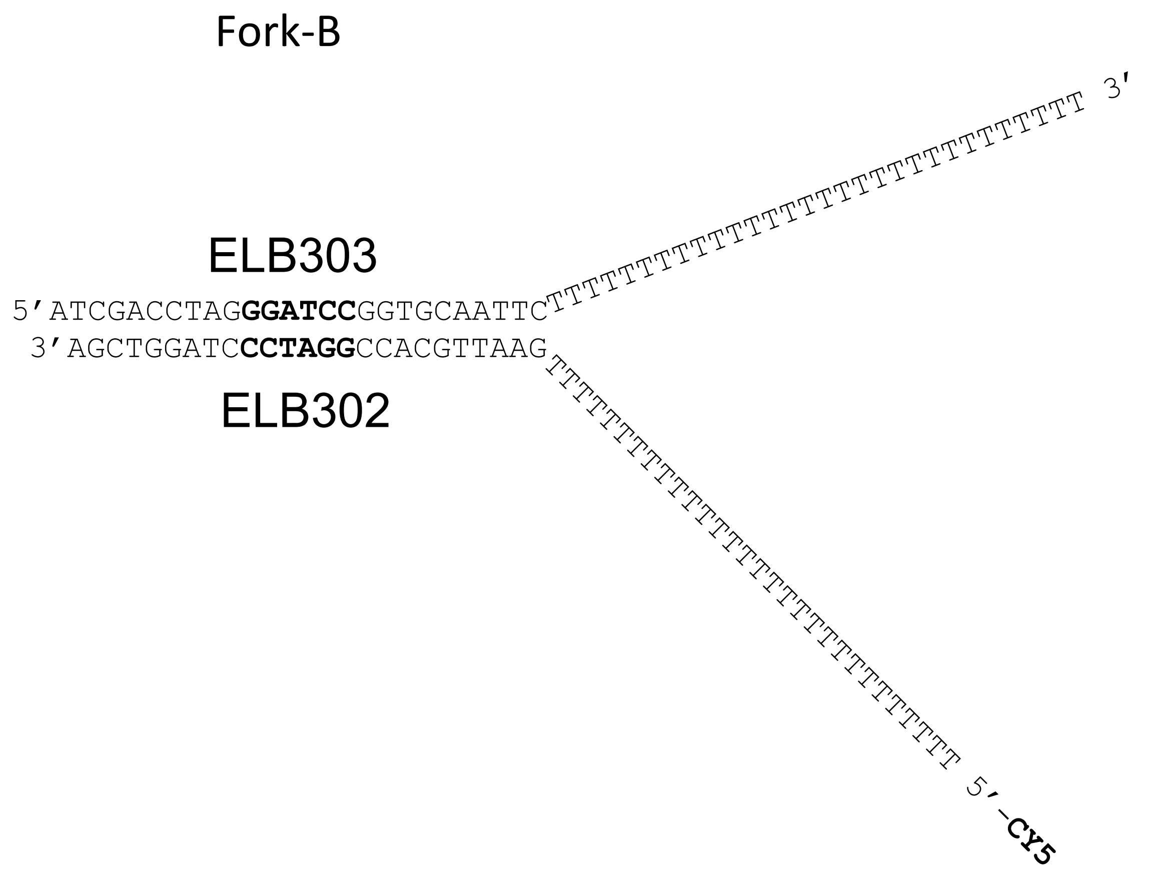

Fork-B
ELB303
5’ATCGACCTAGGGATCCGGTGCAATTC
3’AGCTGGATCCCTAGGCCACGTTAAG
ELB302
3’
5’-CY5

**C.**

**Figure S6**

| Substrate | Name | 5'-3' Sequence |
|---|---|---|
| Fork-A | 5'-Cy5- Mw12 | 5'-GTCGGATCCTCTAGACAGCTCCATGATCACTGGCACTGGTAGAATTCGGC |
| | MW14 | 5'-CAACGTCATAGACGATTACATTGCTACATGGAGCTCTCTAGAGGATCCGA |
| Fork-Me | 5'-Cy5-MePhos-1 | 5'-GTCGGATCCTCTAGACAGC**A**TCCATGATCACTGGCACTGGTAGAATTCGGC |
| | MW14+T | 5'-CAACGTCATAGACGATTACATTGCTACATGGA**T**GCTGTCTAGAGGATCCGA |
| Fork-AP | Abasic-3 | 5'-GTCGGATCCTCTAGACAGCTC**C**ATGATCACTGGCACTGGTAGAATTCGGC |
| | MW14 | 5'-CAACGTCATAGACGATTACATTGCTACATGGAGCTCTCTAGAGGATCCGA |
| Fork-S | 5'-Cy5-Phosphoro2 | 5'-GTCGGATCCTCTAGACAG**CT**CCATGATCACTGGCACTGGTAGAATTCGGC |
| | MW14 | 5'-CAACGTCATAGACGATTACATTGCTACATGGAGCTCTCTAGAGGATCCGA |
| G4 Quadruplex DNA | ADDx3 POLIA 3' | 5'-TCGCCACGTTTCGCCGTTTGCGGGGGTTTCTGCGAGGAACTTTGGAAAAAAAAAAAAAA |

### Figure S7

#### Sequence

The **highlighted portions of sequence** are where there is 75% agreement between all predictors in the database for this region being disordered.

>HelQ
**MDECGSRIRRRVSLPKRN**RPSLGCIFGAP**T**AA**ELVPGD**
**EGKEEEEMVAENRRRKT**AGVLPVEVQPLLLSDSPECLV
LGGGDTNPDLLRHMPTDRGVGDQPN**DSEVDM**FGDYDSF
TENSFIAQVDDLEQKYMQLPEHKKHATDFATENLCSES
IKNKLSITTIGNLTELQ**TD**K**HTENQSGY**EGVTIEPGAD
LLYDVPSSQAIYFENL**QNSSNDLGDHSMKERDWKSSSH**
**NTVNEELPHNCIEQPQQNDESSSKVRTSSDMNRRKSIK**
**DHLKNA**MTGNAKAQTPIFSRSKQLKDTLLSEEINVAKK
TVESSSNDLGPFYSLPSKVRDLYAQFKGIEKLYEWQHT
CLTLNSVQERKNLIYSLPTSGGKTLVAEILMLQELLCC
RKDVLMILPYVAIVQEKISGLSSFGIELGFFVEEYAGS
KGRFPPTKRREKKSLYIATIEKGHSLVNSLIETGRIDS
LGLVVVDELHMIGEGSRGATLEMTLAKILYTSKTTQII
GMSATLNNVEDLQKFLQAEYYTSQFRPVELKEYLKIND
TIYEVDSKAENGMTFSRLLNYKYSDTLKKMDPDHLVAL
VTEVIPNYSCLVFCPSKKNCENVAEMICKFLSKEYLKH
KEKEKCEVIKNLKNIGNGNLCPVLKRTIPFGVAYHHSG
LTSDERKLLEEAYSTGVLCLFTCTSTLAAGVNLPARRV
ILRAPYVAKEFLKRNQYKQMIGRAGRAGIDTIGESILI
LQEKDKQQVLELITKPLENCYSHLVQEFTKGIQTLFLS
LIGLKIATNLDDIYHFMNGTFFGVQQKVLLKEKSLWEI
TVESLRYLTEKGLLQKDTIYKSEEEVQYNFHITKLGRA
SFKGTIDLAYCDILYRDLKKGLEGLVLESLLHLIYLTT
PYDLVSQCNPDWMIYFRQFSQLSPAEQNVAAILGVSES
FIGKKASGQAIGKKVDKNVVNRLYLSFVLYTLLKETNI
WTVSEKFNMPRGYIQNLLTGTASFSSCVLHFCEELEEF
WVYRALLVELTKKLTYCVKAELIPLMEVTGVLEGRAKQ
LYSAGYKSLMHLANANPEVLVRTIDHLSRRQAKQIVSS
AKMLLHEKAEALQEEVEELLRLPSDFP**GAVASST**DK**A**

A.

Figure S3

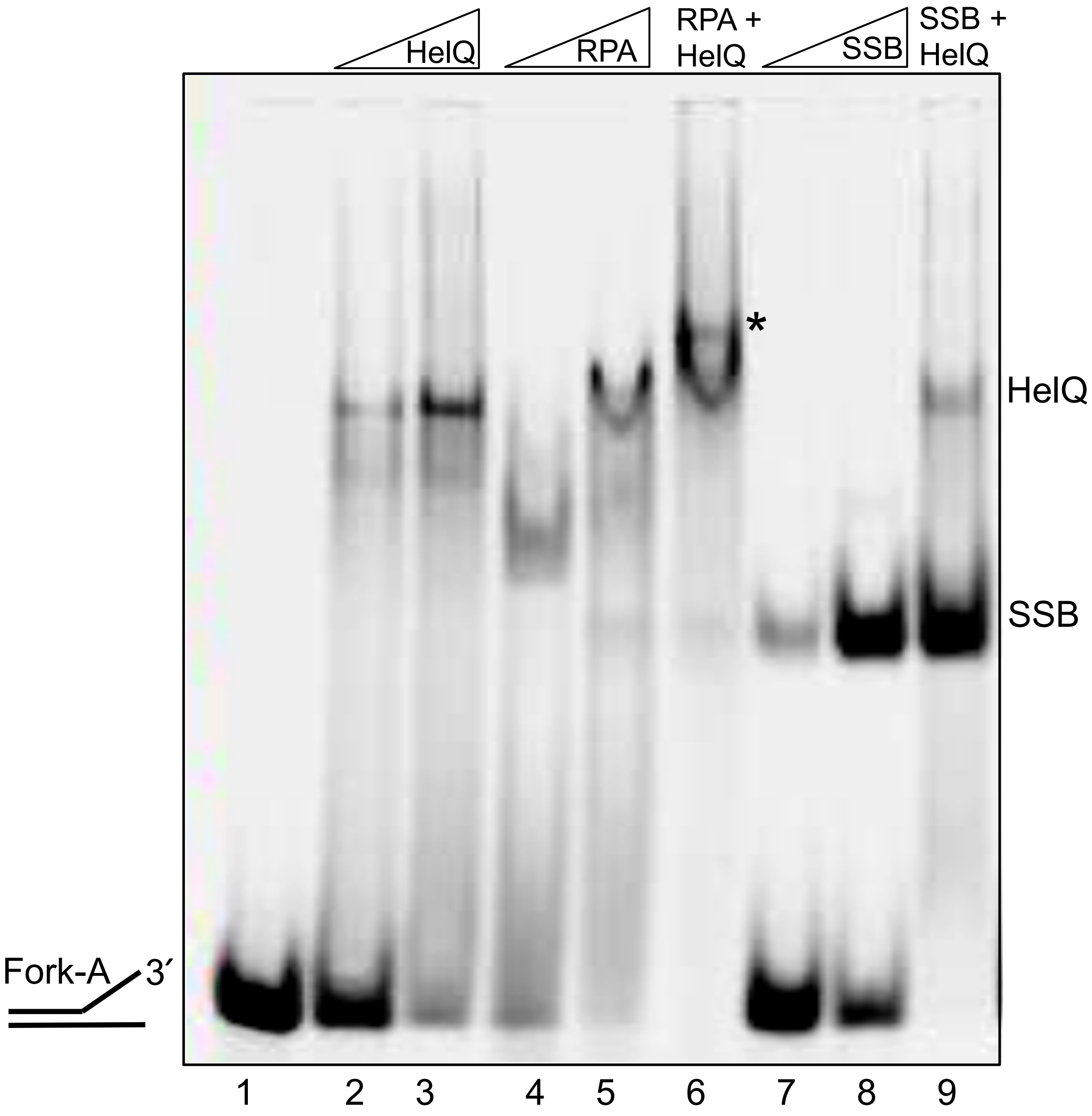

**B.**

**Figure S3**

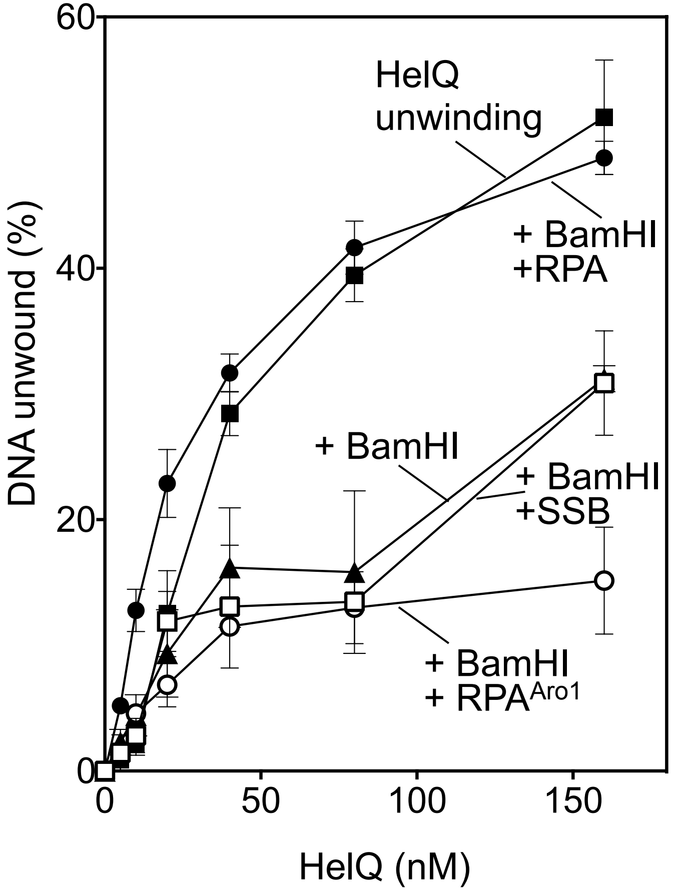
60
40
20
0
0
50
100
150
DNA unwound (%)
HelQ (nM)
HelQ
unwinding
+ BamHI
+RPA
+ BamHI
+ BamHI
+SSB
+ BamHI
+ RPA^Aro1

C.

Figure S3

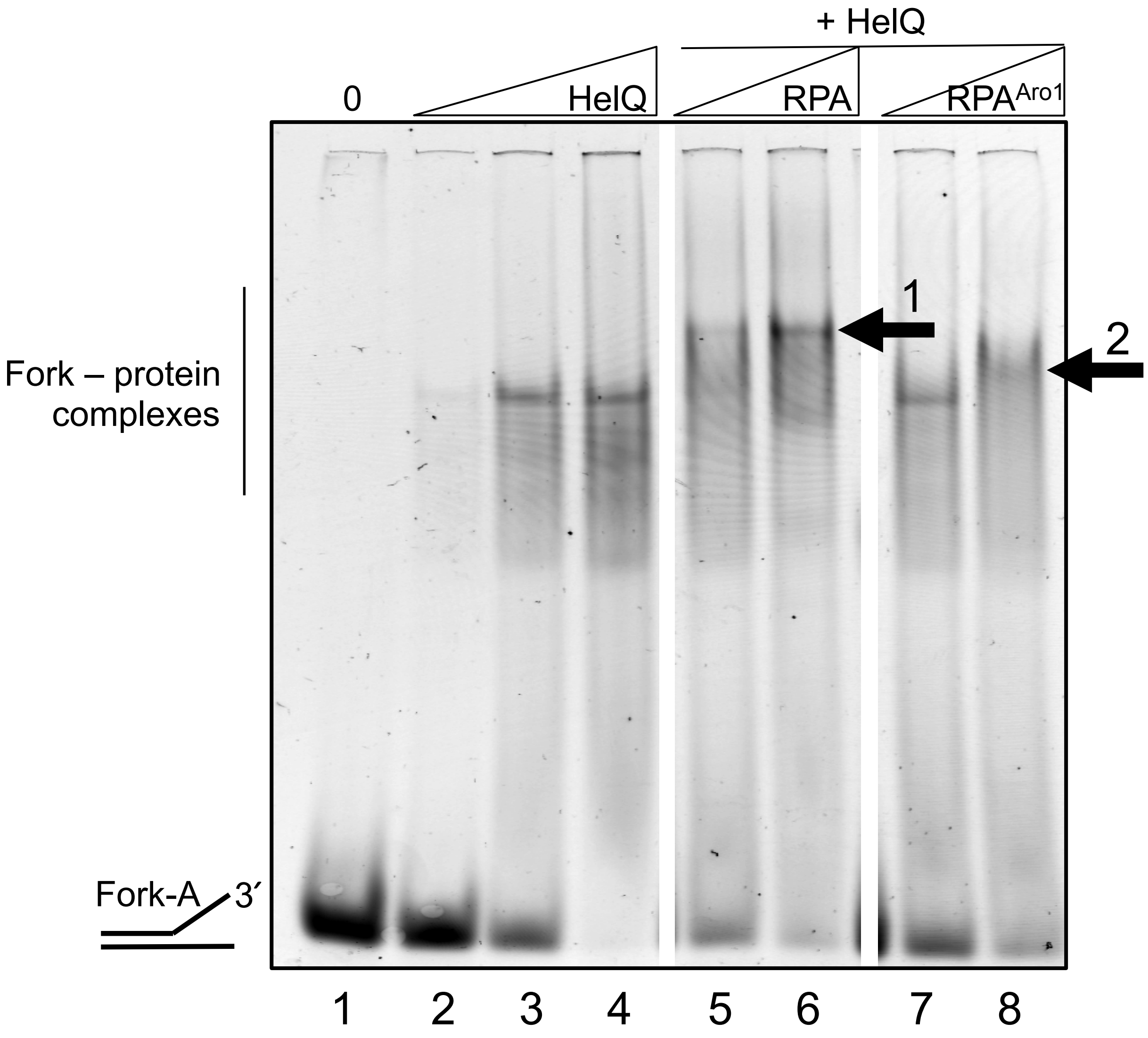
+ HelQ
0
HelQ
RPA
RPA^Aro1
Fork – protein
complexes
1
2
Fork-A
3´
1 2 3 4 5 6 7 8

**Figure S3**

**D.**

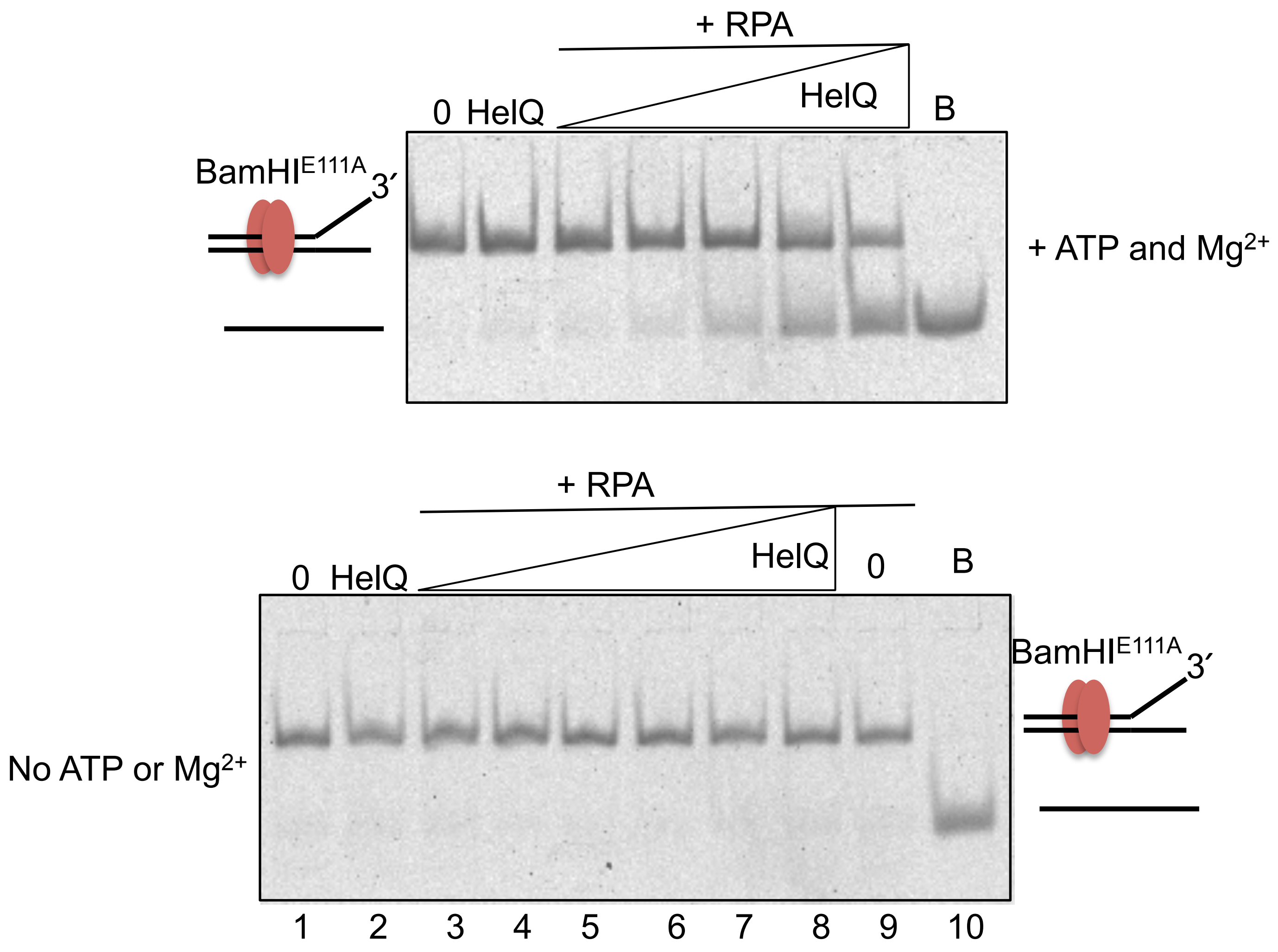

+ RPA
0 HelQ
HelQ
B
BamHI$^{E111A}$ 3′
+ ATP and Mg$^{2+}$
+ RPA
0 HelQ
HelQ
0
B
BamHI$^{E111A}$ 3′
No ATP or Mg$^{2+}$
1 2 3 4 5 6 7 8 9 10

# E.

**Figure S3**

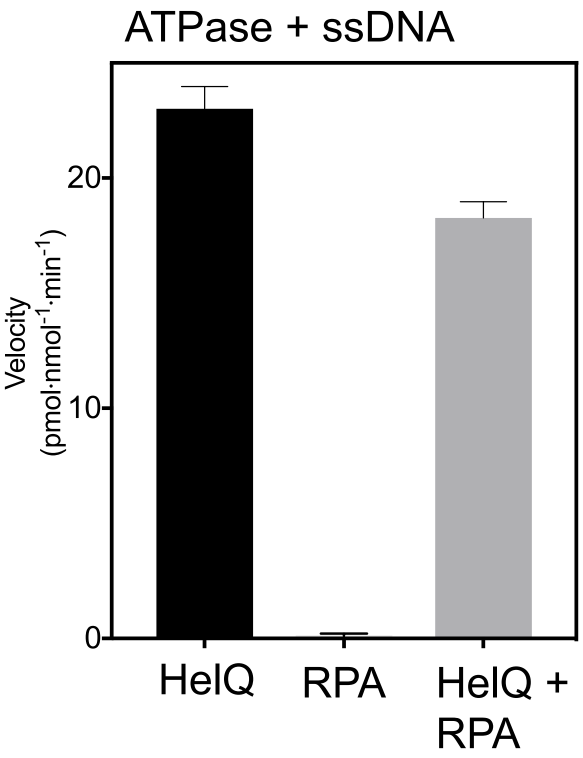

ATPase + ssDNA
Velocity
(pmol·nmol-1·min-1)
20
10
0
HelQ
RPA
HelQ +
RPA

**F.**

**Figure S3**

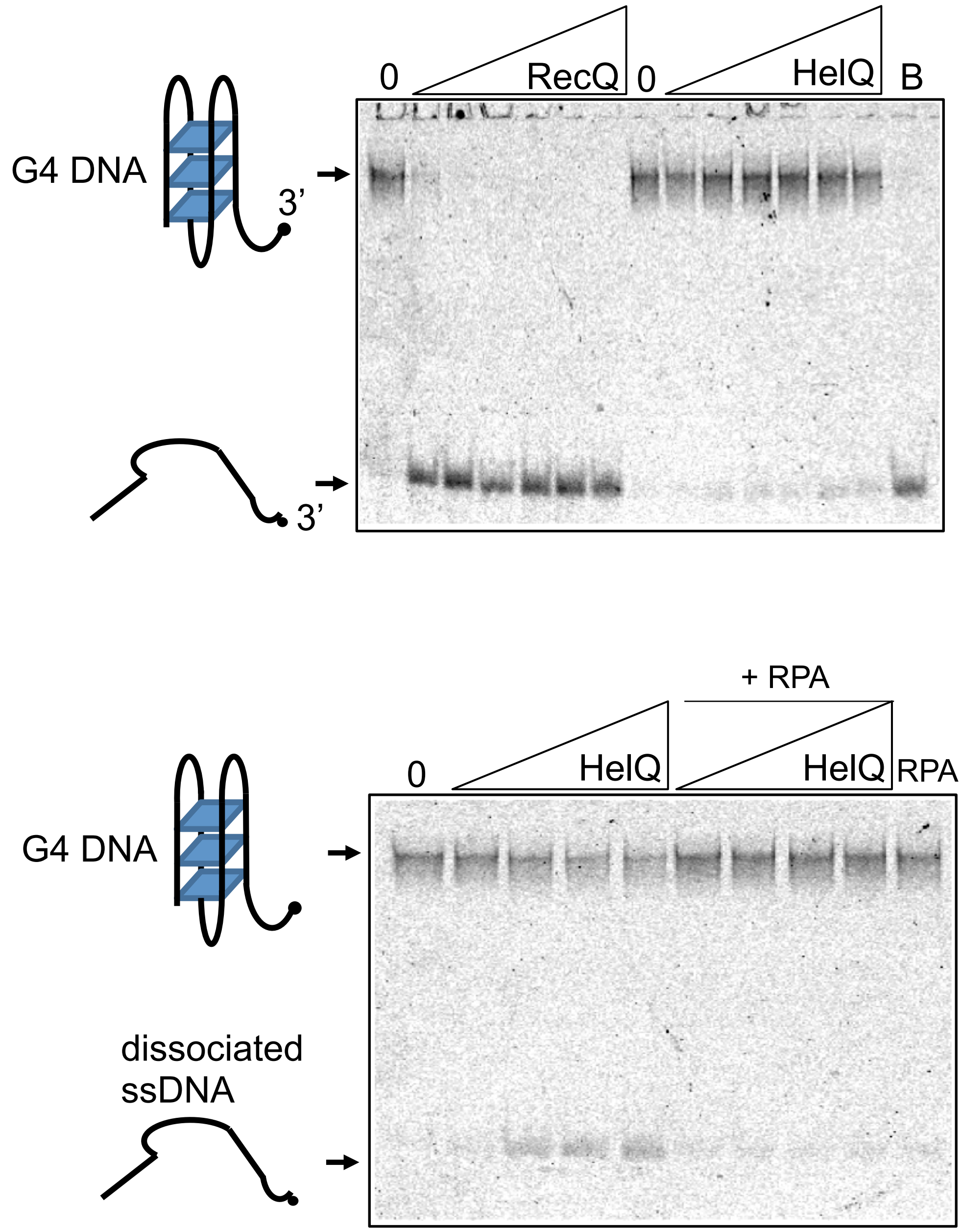
0
RecQ
0
HelQ
B
G4 DNA
3’
3’
+ RPA
0
HelQ
HelQ
RPA
G4 DNA
dissociated
ssDNA

# G.

Figure S3

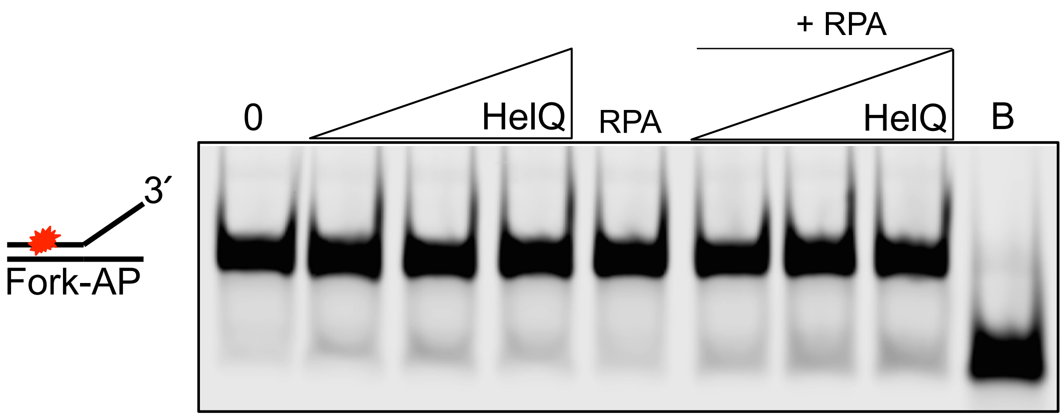

### Figure S4

**A.**

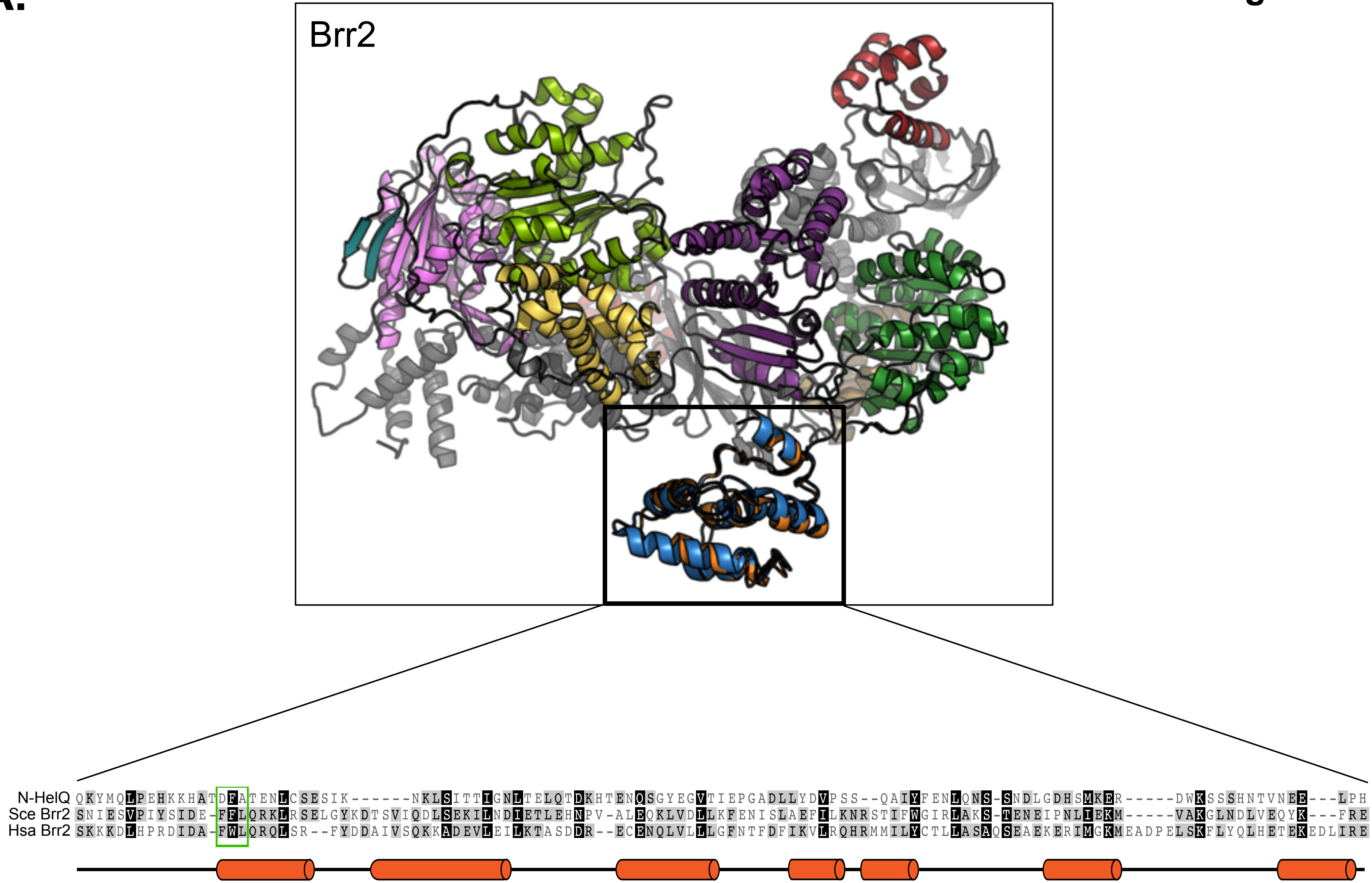

Brr2
N-HelQ QKYMQLPEHKKHATDFATENLCSESIK------NKLSITTIGNLTELQTDKHTENQSGYEGVTIEPGADLLYDVPSS--QAIYFENLQNS-SNDLGDHSMKER-----DWKSSSHNTVNEE---LPH
Sce Brr2 SNIESVPIYSIDE-FFLQRKLRSELGYKDTSVIQDLSEKILNDIETLEHNPV-ALEQKLVDLLKFENISLAEFILKNRSTIFWGIRLAKS-TENEIPNLIEKM-----VAKGLNDLVEQYK---FRE
Hsa Brr2 SKKKDLHPRDIDA-FWLQRQLSR--FYDDAIVSQKKADEVLEILKTASDDR--ECENQLVLLLGFNTFDFIKVLRQHRMMILYCTLLASAQSEAEKERIMGKMEADPELSKFLYQLHETEKEDLIRE

# B.

**Figure S4**

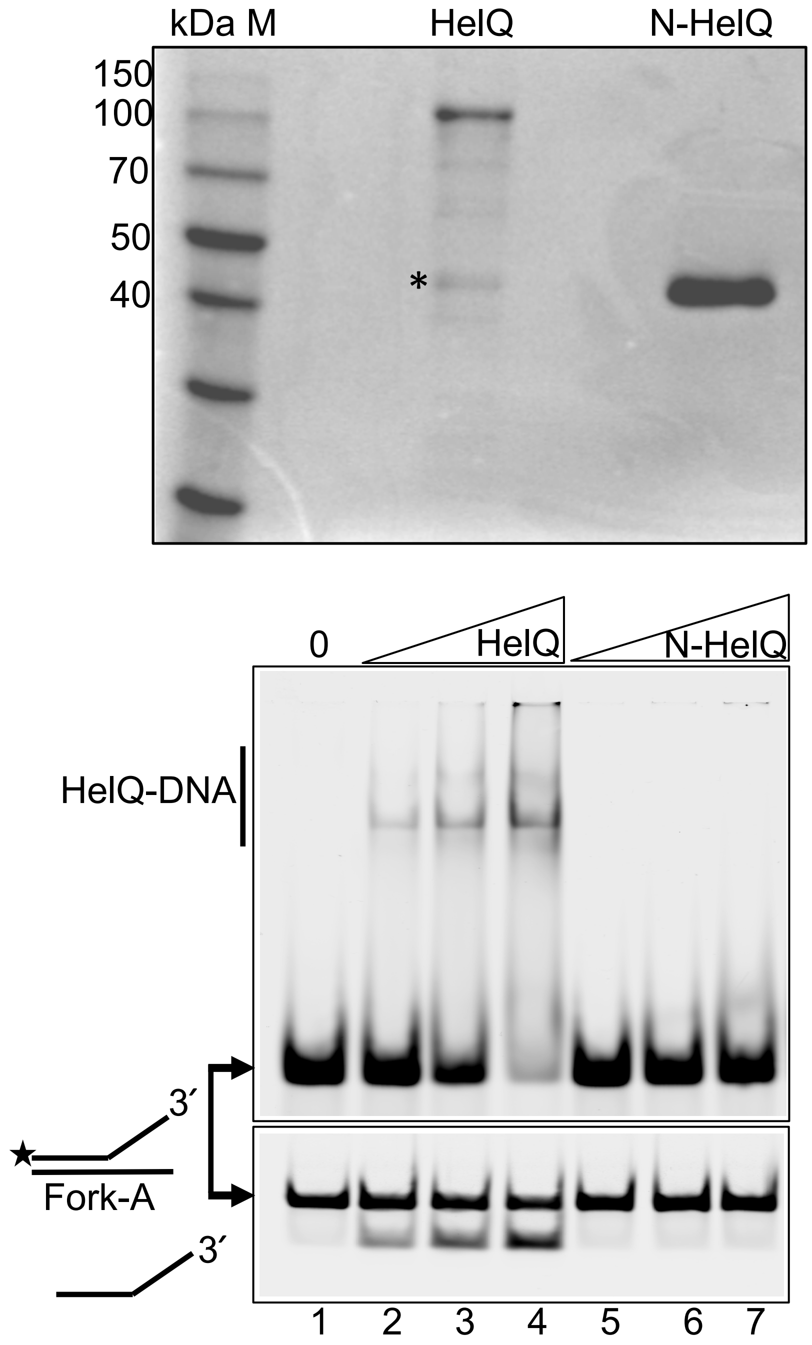
kDa M
HelQ
N-HelQ
150
100
70
50
40
*
0
HelQ
N-HelQ
HelQ-DNA
3′
Fork-A
3′
1
2
3
4
5
6
7

Figure S4

C.

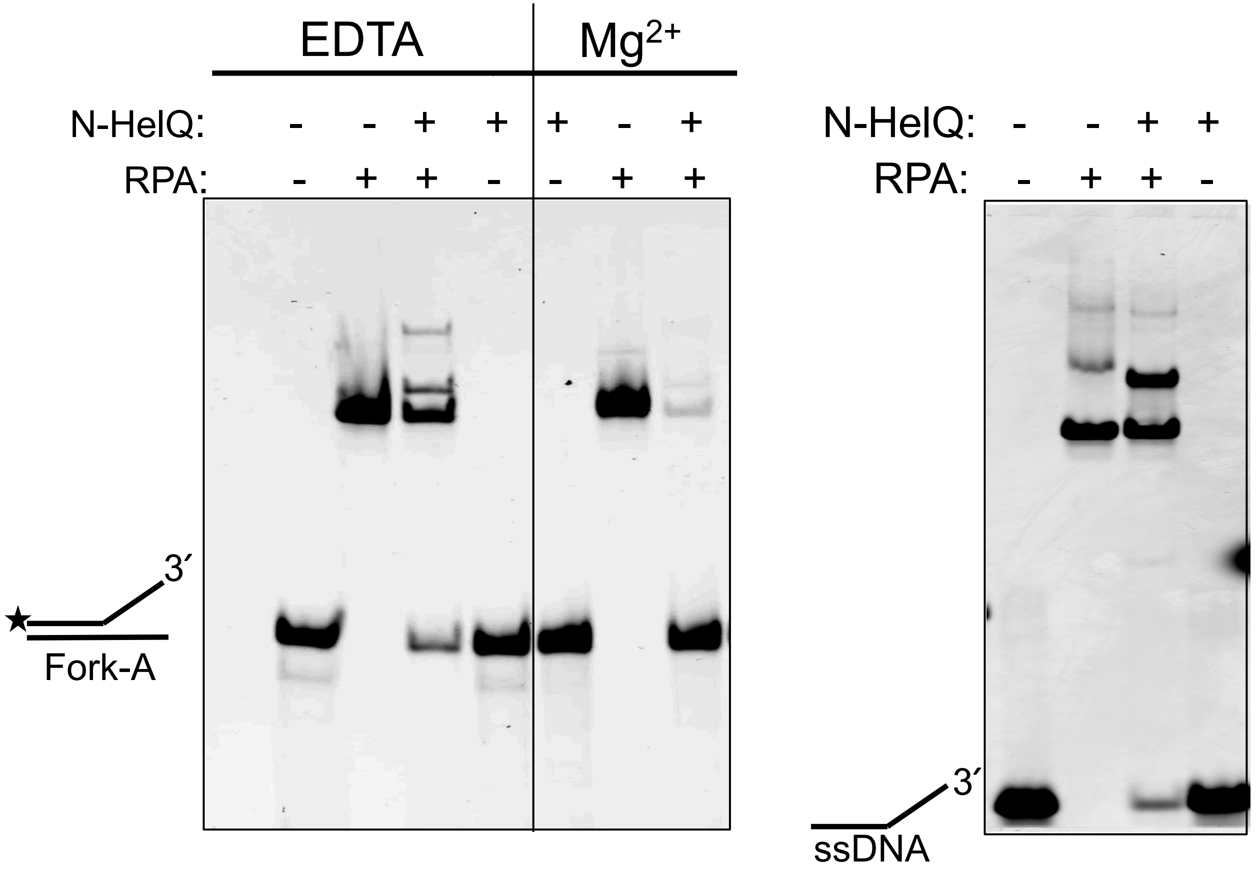

**Figure S5**

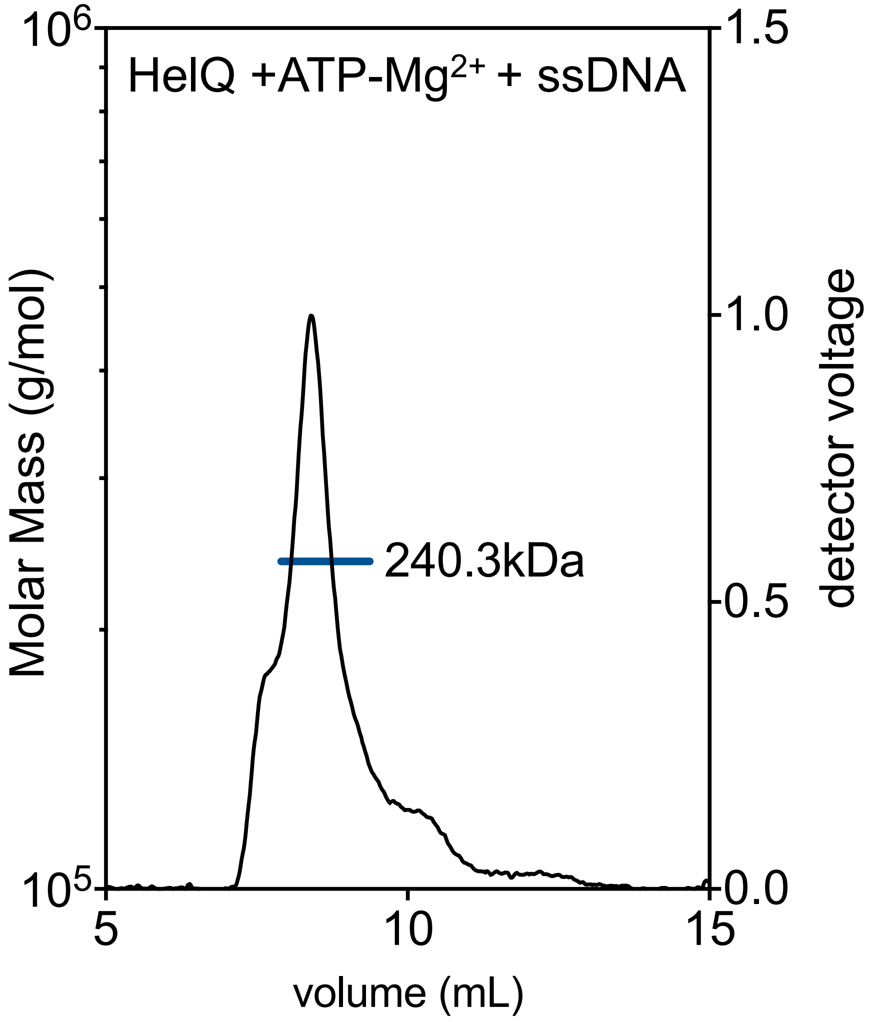

HelQ +ATP-Mg2+ + ssDNA
240.3kDa
Molar Mass (g/mol)
detector voltage
volume (mL)
10^6
10^5
5
10
15
1.5
1.0
0.5
0.0

# A.

**Figure S6**

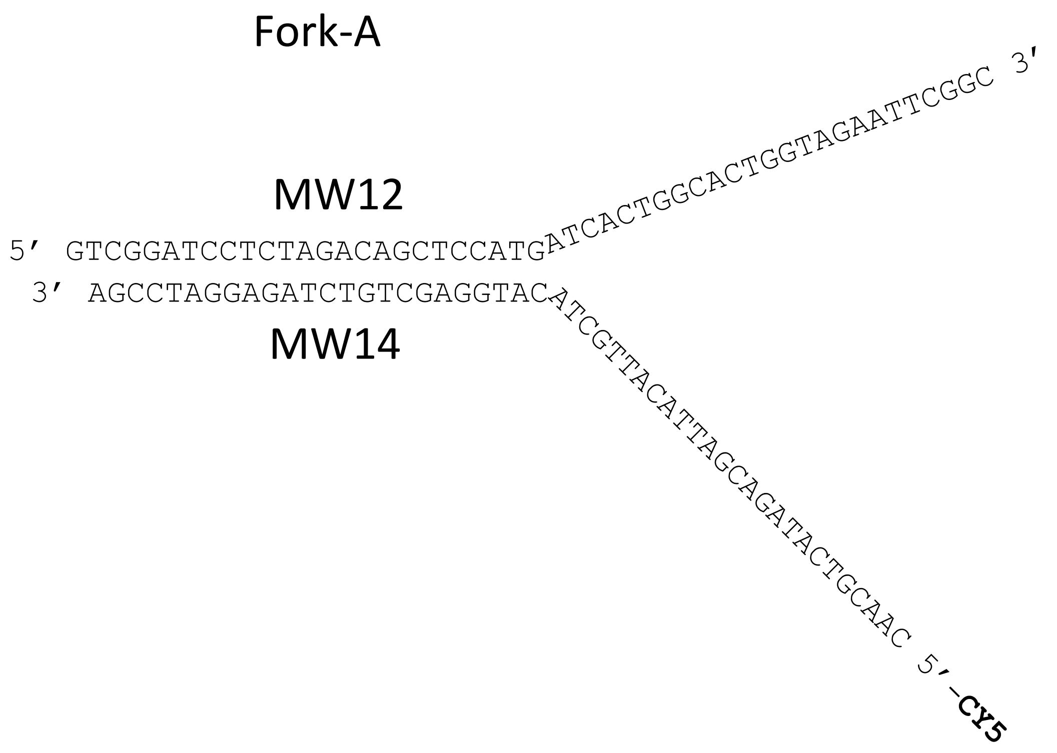
Fork-A
MW12
5′ GTCGGATCCTCTAGACAGCTCCATG
ATCACTGGCACTGGTAGAATTCGGC 3′
3′ AGCCTAGGAGATCTGTCGAGGTAC
ATCGTTACATTAGCAGATACTGCAAC 5′-CY5
MW14

**B.**

**Figure S6**

Fork-B

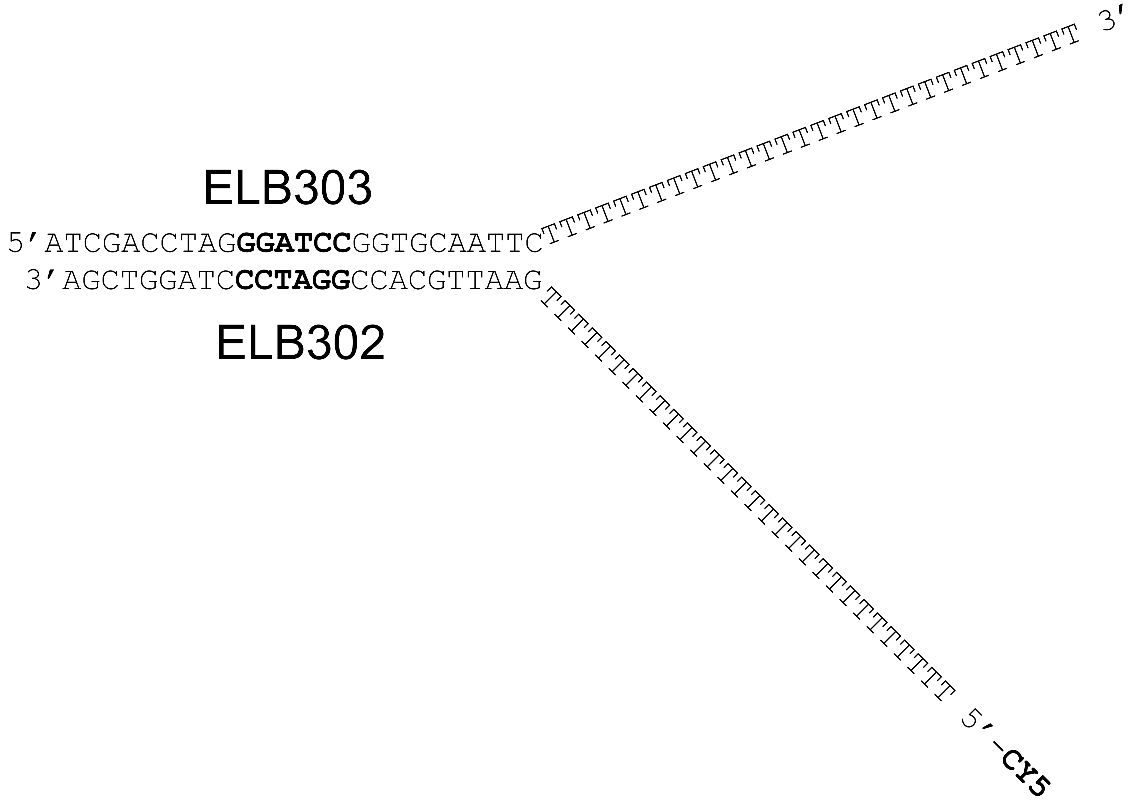

**C.**

**Figure S6**

| Substrate | Name | 5'-3' Sequence |
|---|---|---|
| Fork-A | 5'-Cy5- Mw12 | 5'-GTCGGATCCTCTAGACAGCTCCATGATCACTGGCACTGGTAGAATTCGGC |
| | MW14 | 5'-CAACGTCATAGACGATTACATTGCTACATGGAGCTCTCTAGAGGATCCGA |
| Fork-Me | 5'-Cy5-MePhos-1 | 5'-GTCGGATCCTCTAGACAGC**A**TCCATGATCACTGGCACTGGTAGAATTCGGC |
| | MW14+T | 5'-CAACGTCATAGACGATTACATTGCTACATGGA**T**GCTGTCTAGAGGATCCGA |
| Fork-AP | Abasic-3 | 5'-GTCGGATCCTCTAGACAGCTC**C**ATGATCACTGGCACTGGTAGAATTCGGC |
| | MW14 | 5'-CAACGTCATAGACGATTACATTGCTACATGGAGCTCTCTAGAGGATCCGA |
| Fork-S | 5'-Cy5-Phosphoro2 | 5'-GTCGGATCCTCTAGACAG**CT**CCATGATCACTGGCACTGGTAGAATTCGGC |
| | MW14 | 5'-CAACGTCATAGACGATTACATTGCTACATGGAGCTCTCTAGAGGATCCGA |
| G4 Quadruplex DNA | ADDx3 POLIA 3' | 5'-TCGCCACGTTTCGCCGTTTGCGGGGGTTTCTGCGAGGAACTTTGGAAAAAAAAAAAAAA |

**Figure S7**

#### Sequence

The **highlighted portions of sequence** are where there is 75% agreement between all predictors in the database for this region being disordered.

>HelQ
**MDECGSRIRRRVSLPKRN**RPSLGCIFGAP**T**AA**ELVPGD**
**EGKEEEEMVAENRRRKT**AGVLPVEVQPLLLSDSPECLV
LGGGDTNPDLLRHMPTDRGVGDQPN**DSEVDM**FGDYDSF
TENSFIAQVDDLEQKYMQLPEHKKHATDFATENLCSES
IKNKLSITTIGNLTELQ**TD**K**HTENQSGY**EGVTIEPGAD
LLYDVPSSQAIYFENL**QNSSNDLGDHSMKERDWKSSSH**
**NTVNEELPHNCIEQPQQNDESSSKVRTSSDMNRRKSIK**
**DHLKNA**MTGNAKAQTPIFSRSKQLKDTLLSEEINVAKK
TVESSSNDLGPFYSLPSKVRDLYAQFKGIEKLYEWQHT
CLTLNSVQERKNLIYSLPTSGGKTLVAEILMLQELLCC
RKDVLMILPYVAIVQEKISGLSSFGIELGFFVEEYAGS
KGRFPPTKRREKKSLYIATIEKGHSLVNSLIETGRIDS
LGLVVVDELHMIGEGSRGATLEMTLAKILYTSKTTQII
GMSATLNNVEDLQKFLQAEYYTSQFRPVELKEYLKIND
TIYEVDSKAENGMTFSRLLNYKYSDTLKKMDPDHLVAL
VTEVIPNYSCLVFCPSKKNCENVAEMICKFLSKEYLKH
KEKEKCEVIKNLKNIGNGNLCPVLKRTIPFGVAYHHSG
LTSDERKLLEEAYSTGVLCLFTCTSTLAAGVNLPARRV
ILRAPYVAKEFLKRNQYKQMIGRAGRAGIDTIGESILI
LQEKDKQQVLELITKPLENCYSHLVQEFTKGIQTLFLS
LIGLKIATNLDDIYHFMNGTFFGVQQKVLLKEKSLWEI
TVESLRYLTEKGLLQKDTIYKSEEEVQYNFHITKLGRA
SFKGTIDLAYCDILYRDLKKGLEGLVLESLLHLIYLTT
PYDLVSQCNPDWMIYFRQFSQLSPAEQNVAAILGVSES
FIGKKASGQAIGKKVDKNVVNRLYLSFVLYTLLKETNI
WTVSEKFNMPRGYIQNLLTGTASFSSCVLHFCEELEEF
WVYRALLVELTKKLTYCVKAELIPLMEVTGVLEGRAKQ
LYSAGYKSLMHLANANPEVLVRTIDHLSRRQAKQIVSS
AKMLLHEKAEALQEEVEELLRLPSDFP**GAVASST**DK**A**
